# Context, not contradiction: a pan-cancer framework for conforming and nonconforming co-expression programmes

**DOI:** 10.64898/2026.11.02.763469

**Authors:** Safir Ullah Khan, Ava Storgard, Alfonso Mendez Tenorio, Dwayne G. Stupack

**Author notes:** Correspondence: Dwayne G. Stupack, [ ]; tel. [8588221150].

## Abstract

Reports of a gene’s role in cancer frequently conflict: the same transcript is described as an adverse prognostic marker in one tumour type and a favourable one in another, and such discordance is usually attributed to cohort differences or to irreproducibility. An alternative explanation is that the transcriptional company a gene keeps genuinely differs between lineages. That claim is testable, but testing it requires separating real lineage-dependence from three artefacts that pervade pan-cancer co-expression analysis: public summaries report only coefficients above a fixed threshold, a gene in a recurrently gained region acquires correlates through dosage alone, and conservation across cancer types is assumed rather than measured. We present a framework that addresses all three for any index gene. Correlations are recomputed threshold-free from each cohort’s expression matrix; transcripts are partitioned into cis and trans sets relative to the index gene after retired symbols are resolved; a recurrence rule freezes the gene set before any downstream analysis; cohorts are scored against expression- decile-matched permutation nulls; and copy number is residualised out using the cis set as a built-in positive control, so the adjustment is shown to work before the trans result is read. Applied to PTK2, encoding focal adhesion kinase, across 17 TCGA cohorts, expression was associated with poor outcome in several cancers and better outcome in others, independently of tumour purity. The framework shows this is not noise: cohorts separate into conforming and nonconforming groups enriched for different biology. The 44-transcript trans programme is unchanged by copy-number adjustment while the cis control collapses by 38%, and co- expression is recoverable within individual tumours in 133 of 243 single-cell cases.

## 1. Introduction

Whether a given gene helps or harms in cancer is often reported inconsistently. The same transcript is described as an adverse prognostic marker in one tumour type and a favourable one in another; pathway associations reported in one cohort fail to replicate in the next; and the literature accumulates claims that cannot all be correct. The usual explanations are methodological, invoking cohort composition, platform or statistical treatment, and the usual remedy is a larger meta-analysis. A different explanation is that the discordance is real, and that the transcriptional company a gene keeps differs between tissue lineages, so that the same gene participates in different programmes in different cancers. This is an empirical claim rather than a caveat, and it can be tested. Doing so requires separating genuine lineage-dependence from three artefacts that pervade pan-cancer co-expression analysis.

The first concerns measurement. Much of what is known about any gene’s transcriptional context derives from co-expression summaries returned by public expression portals, which report only genes whose correlation exceeds a fixed coefficient. A gene falling below that threshold in one tumour type is indistinguishable from a gene never measured, and any consensus assembled across tumour types is biased toward whichever cohorts happen to clear it. The second concerns causation. A large fraction of the cancer genome is subject to recurrent arm-level gain or loss, and a gene within such a region correlates with its chromosomal neighbours for a purely mechanical reason, since amplification raises both together. Unless copy number is controlled, a co-expression list for such a gene is in part a map of its amplicon. The third concerns conservation. Co-expression results are frequently pooled across cohorts, or reported for one cohort and generalised, which presumes the programme is the same everywhere. Whether a gene’s transcriptional context is conserved across cancer types is an empirical question that pooling makes unanswerable. Addressing all three requires recomputing correlations without a threshold, separating dosage from regulation in a way that can be checked rather than asserted, and testing conservation rather than assuming it. The second is hardest, because an adjustment that removes nothing is indistinguishable from one that was never needed. We therefore exploit a feature available for any gene in a recurrently altered region: its own chromosomal neighbours. Transcripts near the index gene must lose signal when copy number is removed, since for them dosage is the true explanation, and they therefore furnish a positive control built into the analysis itself.

We demonstrate the framework on PTK2, which encodes focal adhesion kinase (FAK), a cytoplasmic tyrosine kinase central to cell adhesion, movement and survival and an active target for therapeutic intervention [1]. PTK2 is a demanding test case, because all three problems apply to it acutely. It lies at 8q24.3, telomeric to MYC, within one of the most frequently gained regions of the cancer genome, so elevated FAK may reflect regional gain as readily as selection on FAK itself. Much of what is reported about its transcriptional context derives from portal co-expression summaries [2,3]. And FAK signalling is broadly assumed to operate identically across cancers, though this has rarely been examined: FAK is held to promote motility, invasion and epithelial-to-mesenchymal transition, support cancer stem cell properties, modify the tumour microenvironment, suppress anti-tumour immunity and sustain vascularization, while activating pro-survival pathways and integrating integrin signalling with adhesion [4–7]. Recent work in ovarian cancer illustrates both arms: inhibiting tumour FAK releases omega-3 fatty acid-containing exosomes that program CXCL13-producing peritoneal macrophages [8], while tumour FAK sustains fibrillar collagen expression in ascites tumour cells [9].

A previous pan-cancer survey of PTK2 across 33 TCGA cohorts reported survival associations, pathway enrichment and immune correlates from bulk expression [10]. That analysis established the breadth of the question but did not address any of the three artefacts above: correlations were not re-derived without a threshold, copy number was not controlled despite the position of PTK2 on 8q, and no single-cell data were used, so compartment mixing in bulk tissue could not be excluded. The present work differs in treating those three as the obstacles to be removed rather than as caveats.

PTK2 also exhibits precisely the discordance described above: its expression is associated with poor overall survival in several cohorts and better survival in others, a pattern surviving adjustment for tumour purity and so not explained by malignant cell content. Applying the framework across 17 TCGA cohorts, we find this reversal accompanies a real difference in transcriptional context: tumour types separate into conforming and nonconforming groups differing in associated biology rather than in degree, and the programme distinguishing them is not explained by amplicon dosage.

## 2. Materials and Methods

### 2.1. Framework overview

The analysis proceeds in five steps, none of which depends on the identity of the index gene. (i) Correlations between the index gene and every measured transcript are recomputed from each cohort’s expression matrix with no threshold applied. (ii) Transcripts are partitioned into a cis set on the index gene’s own chromosome arm and a trans set elsewhere, after retired and alias symbols are resolved so that neither set contaminates the other. (iii) A recurrence rule specified in advance freezes the gene set before any downstream analysis. (iv) Each cohort is scored against expression-decile-matched permutation nulls drawn within that cohort. (v) Copy number is residualised out, with the cis set as a positive control: the adjustment must attenuate the cis set, and the degree to which it does calibrates how the trans result should be read. An optional sixth step asks whether co-expression is recoverable within individual tumours in single-cell data. Parameter-level detail is given in Supplementary Methods and every fixed parameter in Table S2.

### 2.2. Data sources

Genetic alteration data for 32 TCGA cancers came from cBioPortal [11,12]; expression across tissue state and stage from TNMplot [13], GEO [14,15] and TISIDB [16]; protein from CPTAC through UALCAN [17]; and representative immunohistochemistry from the Human Protein Atlas [18]. Transcript-level expression for the co-expression analyses was taken from the TCGA PanCanAtlas batch-corrected RSEM matrix through UCSC Xena [19], overall survival from the Pan-Cancer Clinical Data Resource [20], tumour purity from ABSOLUTE consensus estimates [21], which derive from SNP-array copy number and are therefore independent of expression, and arm-level copy number and aneuploidy score from the PanCanAtlas aneuploidy resource [22]. Cohorts, sample sizes and software versions are listed in Table S1 and Supplementary Methods.

### 2.3. Step 1: gene symbol resolution

The TCGA matrix carries legacy HUGO symbols, so many transcripts fail to resolve against current records. Retired and alias symbols were mapped using the HGNC complete set [23] through the prev_symbol and alias_symbol fields, previous symbols taking precedence, and only where the target was unambiguous and not already present as a separate row. Chromosomal assignment followed resolution, and unresolvable transcripts were excluded from the partition rather than defaulting to either class. This recovered 66 chromosome 8q transcripts that would otherwise have been misassigned as trans (Table S13).

### 2.4. Step 2: threshold-free re-derivation and cis/trans partition

Spearman correlations between PTK2 and every transcript were computed within each of 19 cohorts with Benjamini-Hochberg false discovery rate control applied within cohort [24], retaining all coefficients regardless of magnitude (Table S5). A transcript entered the consensus where it reached rho at least 0.30 at FDR below 0.05 in at least six of the seventeen cohorts containing at least 100 patients, each cohort contributing at most its 500 strongest correlations per class so that a noisier transcriptome could not dominate. Transcripts resolving to chromosome 8q were classified as cis and all others as trans (Tables S2, S3).

### 2.5. Step 3: cohort scoring against expression-matched nulls

Each cohort was scored by the mean Spearman correlation between PTK2 and the frozen gene set and compared against 1,000 random sets matched on expression decile, so that the null controls for the tendency of highly expressed genes to correlate more readily. Because 1,000 permutations place a floor of 0.001 on the empirical p value, cohorts are compared by z-score. Tiers were assigned by tertile (Table S4).

### 2.6. Step 4: copy-number residualisation with a built-in positive control

Arm-level 8q copy number and the genome-wide aneuploidy score were matched to expression by sample barcode. Partial Spearman correlations were computed by rank-transforming expression and covariates, residualising both PTK2 and each transcript on the covariate matrix by least squares, and correlating the residuals; results in the text refer to the joint adjustment. Chromosome 8q transcripts serve as the positive control, since they must attenuate if the covariate is effective (Table S6).

### 2.7. Step 5: within-tumour validation in single-cell data

Single-cell data came from 38 published studies deposited in the Gene Expression Omnibus [14,15], ArrayExpress and BioProject (accessions in the Data availability statement), restricted to malignant tissue from the ten tumour types with a matching bulk cohort. Within each tumour, cells were aggregated into metacells by k-means clustering on the first 30 principal components of the 2,000 most variable genes, and Spearman correlations between PTK2 and every transcript detected in at least 10% of cells were computed across metacells, each tumour being compared against 1,000 expression-decile-matched random sets drawn within it. PTK2 detection rate and its coefficient of variation across metacells were recorded so that an absence of coupling could be distinguished from an absence of measurable variation (Table S16).

### 2.8. Downstream characterisation of the frozen gene set

Chromosome-arm distribution of the trans transcripts was tested by permutation against the universe of measured transcripts rather than the whole genome, controlling for gene density (Table S7). Pathway analysis used preranked gene set enrichment [25] rather than over- representation testing, which requires a threshold and discards the ranking; each cohort was ranked by correlation with PTK2 and tested against Gene Ontology Biological Process and MSigDB Hallmark [26] using gseapy [27]. Tier contrasts used only terms tested in at least five cohorts of both tiers, averaging only over cohorts in which a term was returned (Tables S8- S10). Transcription factor enrichment used the Enrichr TF Perturbations library [28–30] against the measured universe as background (Tables S11, S12).

### 2.9. Statistical analysis

RNA-Seq and GeneChip comparisons used Student’s t-test at p < 0.05 and stage analysis Spearman’s rank correlation. Survival analysis divided patients at the cohort median of PTK2 expression, comparing groups by log-rank test and estimating hazard ratios by Cox regression fitted on the same binary split, so that the hazard ratio and the log-rank p describe a single comparison; purity-adjusted models added ABSOLUTE purity as a continuous covariate (Table S15). Analyses used Python with pandas, numpy, scipy, statsmodels and gseapy. The final analysis code will be made available at the GitHub repository of S.U.K. (https://github.com/Safir-Ullah-Khan) upon acceptance.

### 2.10. Applying the framework to another index gene

The framework requires only an expression matrix per cohort, the chromosomal location of the index gene, and copy-number calls for the same patients. Three parameters govern the gene set, namely the correlation and false discovery rate thresholds, the number of cohorts in which a transcript must recur, and the number retained per class; the values used here are recorded in Table S2 so that they can be varied. Not every gene is suitable. The analysis is most informative where coding variation is rare, so that expression rather than mutation is the relevant axis; where the gene lies in a region of recurrent copy-number change, so that the cis control is populated; where transcript tracks protein, so that mRNA is a defensible readout; and where existing reports disagree, so that there is a discordance to explain. For a gene in a copy-number- stable region the copy-number step should be omitted rather than reported as a negative result.

## 3. Results

Sections 3.1 to 3.5 establish PTK2 as a test case and define the problem the framework has to solve. A gene is informative for this analysis when four conditions hold: its activity is not dominated by point mutation, so that coding variation can be set aside; it lies in a region of recurrent copy-number change, so that the dosage confounder is present and the cis control is populated; its transcript level tracks both its copy number and its protein, so that mRNA is a defensible readout; and its reported associations with outcome disagree between cohorts, so that there is a discordance to explain. PTK2 satisfies all four, which makes it a demanding rather than a convenient demonstration.

### 3.1. FAK activation via single nucleotide variation is rare

Oncogenes are commonly associated with point mutations that increase activity and are positively selected during tumorigenesis, as distinct from variants of unknown significance, which include possible drivers alongside an abundance of passenger and inactivating mutations [31]. Surveying 19 cancers and more than 10,000 patients, mutations in FAK were far less frequent than in widely expressed oncogenic drivers from COSMIC [32], including receptor tyrosine kinases, GTPases and cytosolic kinases (Figure 1A), and all were categorised as variants of unknown significance.

**Figure 1.**
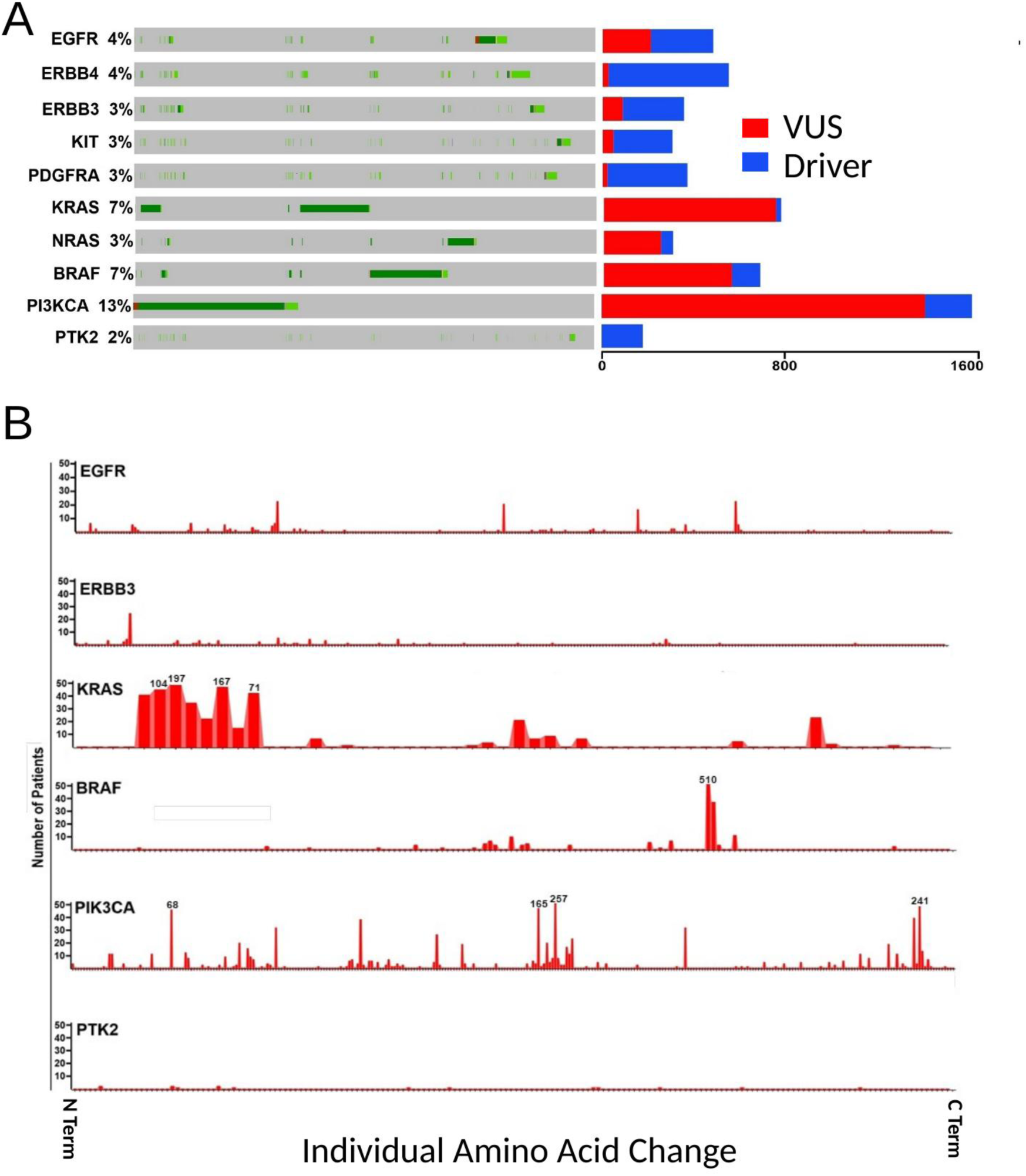
Single nucleotide variation in FAK. (A) Oncoprint profiling of SNVs found in representative oncogenic drivers or in PTK2/FAK across 32 cancers from The Cancer Genome Atlas. The total numbers of variants of characterized drivers or variants of unknown significance (VUS) are shown on the right side of the panel. Mutations in FAK/PTK2 are less frequent than those in key oncogenic drivers like receptor tyrosine kinases and GTPases. (B) Lollipop-style plots mapping SNV/INDEL mutations along the primary amino acid sequence of EGFR, ERBB3, KRAS, BRAF, PIK3CA, and PTK2 based on data in (A). Where a bar exceeds 50 events for a particular site, the total number of events in the database is listed in text above the 50 site. *Alt text: Oncoprint and mutation-frequency panels comparing single nucleotide variants in PTK2 with those in established oncogenic drivers across 32 TCGA cancer types*.

This could reflect incomplete annotation of activity-promoting mutations, since changes in the FERM domain or kinase-adjacent regions produce an open conformation that lowers the threshold for FAK kinase activity [33–40]. We nonetheless found no enrichment for mutations at sites regulating kinase activity, nor in any other region of PTK2 (Figure 1B), whereas other oncogenes show clear enrichment at sites known to promote tumorigenic activity. The low mutation incidence across a large gene, together with the broad and unfocused distribution, is inconsistent with selection for constitutive catalytic activity.

### 3.2. The PTK2 gene is very frequently gained in tumours

Gains or amplifications of PTK2 called by GISTIC were frequent, with ten cancer types carrying an additional copy in at least half of tumours sequenced (Figure 2A). Since tumours are typically aneuploid, we asked whether this simply reflected gains of chromosome 8. Copy number along individual chromosomes, visualised using CAIRN [41] (Figure 2B, Figure S1), showed that PTK2 gains were associated with 8q regional gains rather than whole-chromosome events. Loss of 8q, or whole gains or losses of chromosome 8, were rare except in glioblastoma, where complete loss of PTK2 or gain of the entire chromosome was often observed.

**Figure 2.**
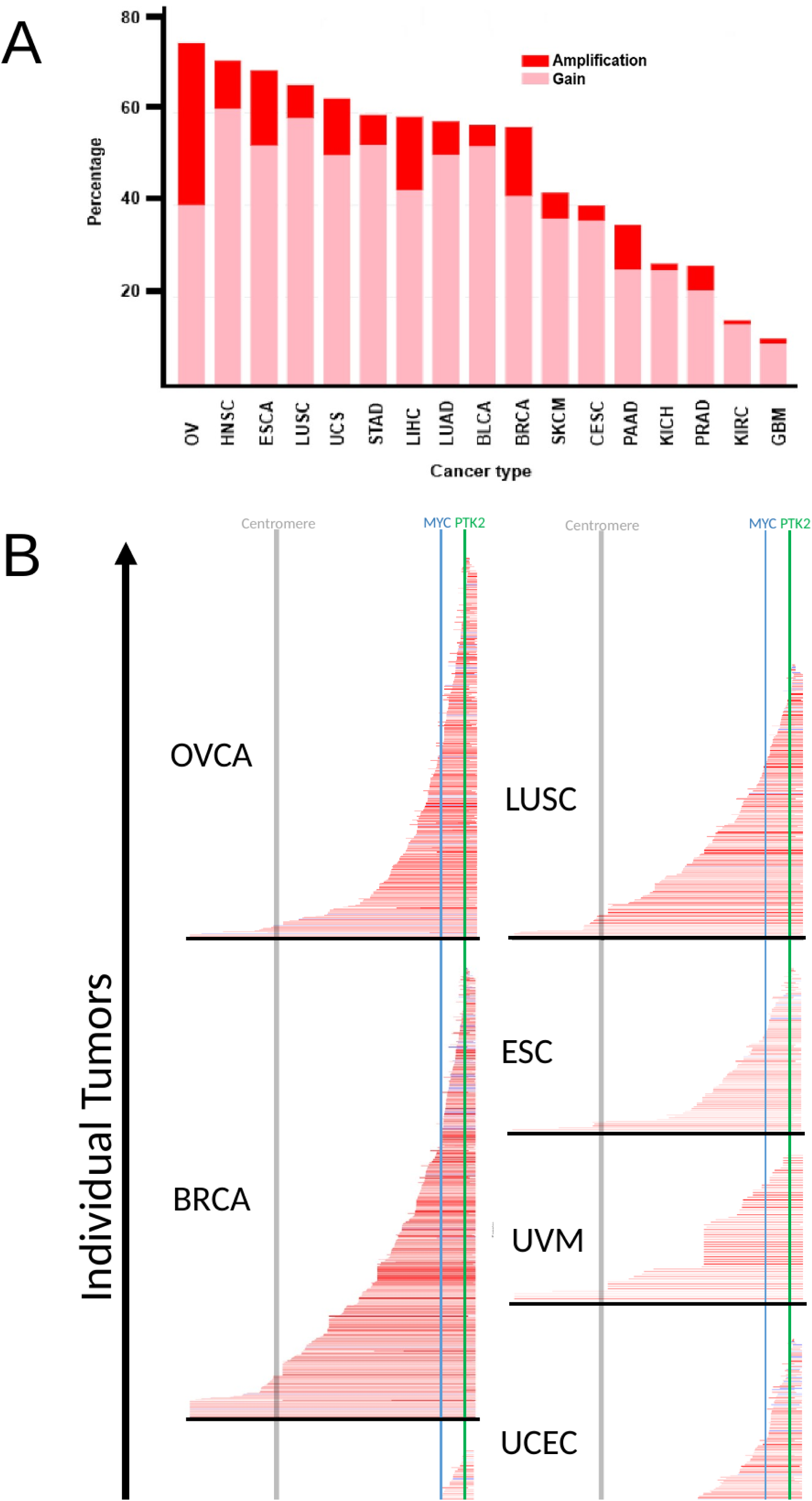
Copy number gains in PTK2/FAK by cancer type. (A) Stacked bar chart highlighting the gain (pink) or amplification (red) of the PTK2 gene across common cancers including ovarian, head and neck, esophageal, lung squamous cell, uterine sarcoma and liver. (B) CAIRN-based representation of chromosomal gains associated with PTK2 amplification for ovarian, breast, cervical, lung squamous cell, esophageal and uterine carcinomas, as well as uveal melanoma, bladder cancer and glioblastoma. The chromosome 8 key at the bottom of each panel denotes the location of PTK2 (FAK) (green line) and MYC (blue line), as well as the centromere (grey line). *Alt text: Stacked bar chart of PTK2 copy number gain and amplification frequency by cancer type, with chromosome 8 copy-number maps for individual tumours*.

Increased PTK2 copies could be collateral, reflecting co-amplification with a neighbouring driver. We therefore examined co-occurrence with MYC, which lies in the same 8q24 region roughly 15 centimorgans away but is not immediately adjacent, so the interval is open to breakage and to selective amplification. Between 20% and 40% of 8q PTK2 gains exclude MYC, depending on tumour type (Figure 2B), arguing that gains of the two genes are selected independently even where collaborative action may occur. The gains are therefore consistent with selection for a biological activity of FAK rather than with passenger status.

### 3.3. PTK2 copy number predicts FAK mRNA expression

Patients with ovarian cancer express FAK, but not MYC, transcripts in proportion to gene copy number [42], and we found this concordance extended across bladder, breast, esophageal, uterine, sarcoma, uveal melanoma and cervical cancer (Figure 3A), indicating that PTK2 copy- number change broadly reflects transcript level. Transcripts were also somewhat elevated in tumour relative to normal tissue (Figure 3B), with exceptions in blood, brain and testis. No broad trend was evident for metastases relative to primary tumours (Figure 3C; Figure S2A), nor overall for tumour stage, though expression trended upwards in colon and kidney malignancy (Figure S2B).

**Figure 3.**
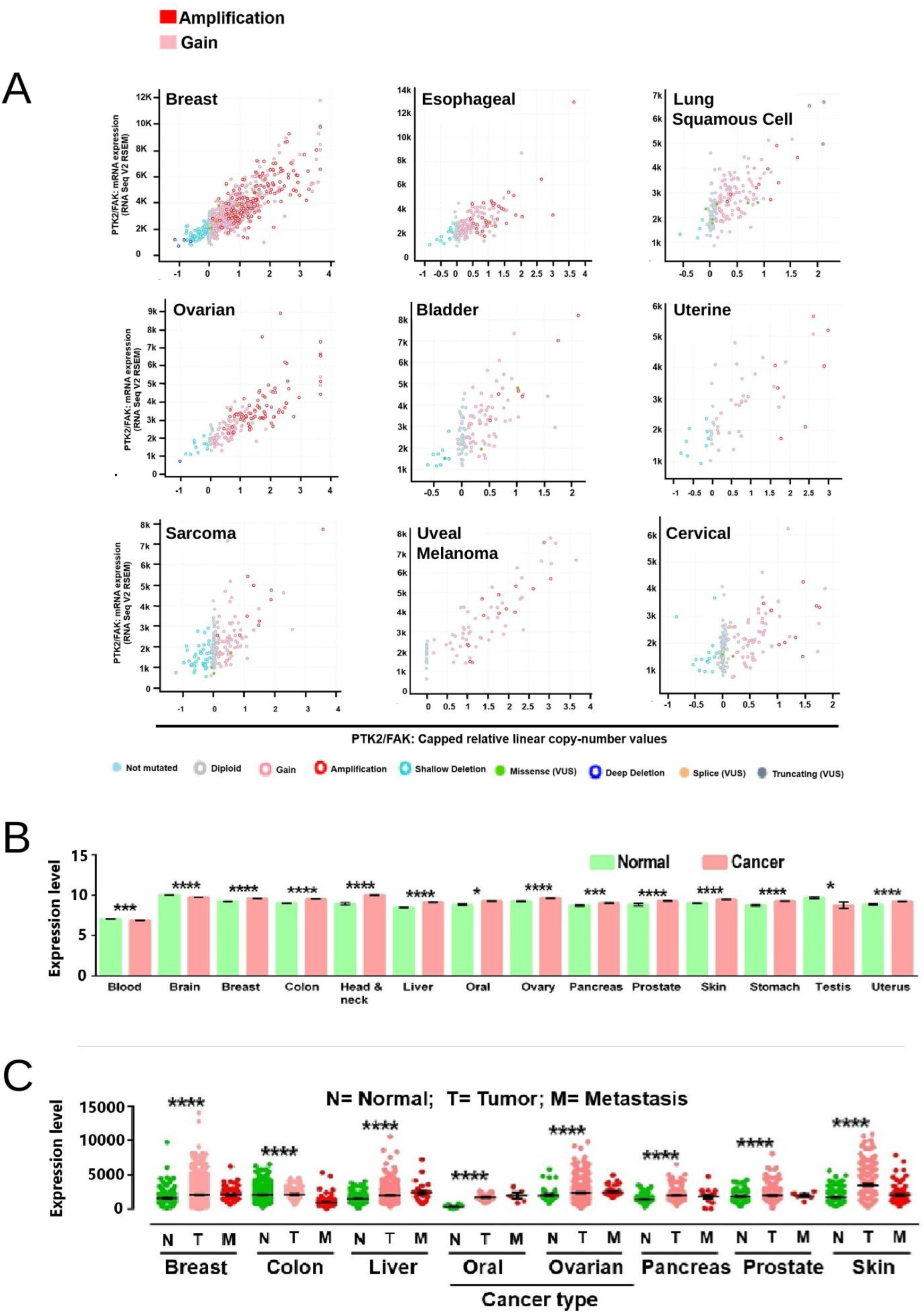
PTK2/FAK mRNA level is elevated in disease and concordant with genomic copy number variation across cancer types. (A) Scatter plots showing the correlation between PTK2/FAK transcript expression levels and gains or losses of the PTK2 gene (GISTIC). (B) Comparison of the relative mRNA levels for PTK2 in normal versus neoplastic tissues. (C) Evaluation of the relative PTK2/FAK expression stratified by primary tumour (T) or metastases (M) versus normal tissues. The p value is as shown (*, p < 0.05; ***, p < 0.01; ****, p < 0.001). No statistical significance was reached between tumour and metastases for any tumour. *Alt text: Scatter plots and box plots relating PTK2 transcript level to PTK2 gene copy number and to tissue state across cancer types*.

### 3.4. FAK protein expression is elevated across most solid tumours

Because protein is not always concordant with mRNA, we evaluated FAK protein using CPTAC datasets. Protein was upregulated in breast, ovarian, pancreatic, head and neck and liver cancer (right panels, Figure 4A-E) and downregulated in glioblastoma (Figure 4F), consistent with glioblastoma showing the lowest frequency of PTK2 gain of any cohort (Figure 2A). Qualitative immunohistochemistry agreed (left panels, Figure 4A-F). These results support a model in which increased PTK2 copy number raises transcript and in turn protein, and establish that transcript is a defensible readout of FAK abundance.

**Figure 4.**
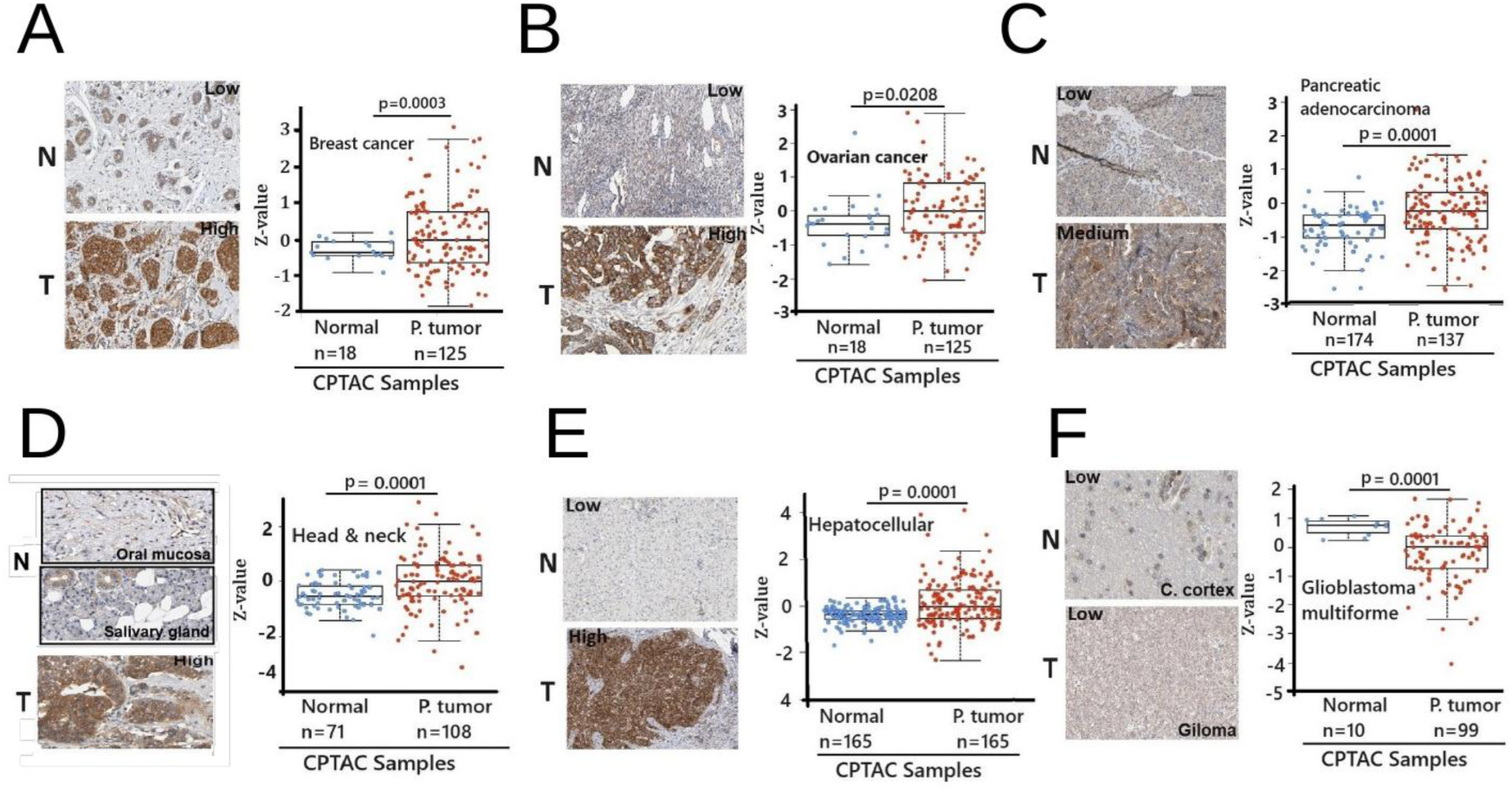
FAK protein expression is elevated across six cancer types. The expression of the FAK protein was evaluated across six cancer types with available CPTAC data. Each panel (A-F) shows representative immunohistochemistry for PTK2 (left) and CPTAC data quantifying FAK expression within tumour types. Tumours shown are breast (A), ovarian (B), pancreatic (C), head and neck (D) and hepatocellular (E) carcinomas, and glioblastoma (F). *Alt text: Six panels pairing immunohistochemistry images for FAK with CPTAC protein quantification in tumour and normal tissue*.

### 3.5. PTK2 association with survival reverses direction between cohorts

Since FAK mRNA did not correlate strongly with metastasis (Figure 3C), we assessed its association with overall survival in TCGA (Figure 5A; Table S15). Outcomes varied by cancer type rather than converging on one direction. Elevated FAK mRNA was an unfavourable prognostic marker in breast cancer (HR 1.87, 95% CI 1.40-2.51, log-rank p < 0.0001), cervical cancer (HR 1.71, 1.07-2.74, p = 0.022), uterine carcinosarcoma (HR 2.02, 1.01-4.04, p = 0.043) and uterine corpus endometrial carcinoma (HR 1.53, 1.02-2.31, p = 0.040), trended in the same direction in sarcoma (HR 1.41, 0.94-2.09, p = 0.091), and was unassociated with outcome in bladder cancer (HR 0.83, 0.63-1.11, p = 0.210).

**Figure 5.**
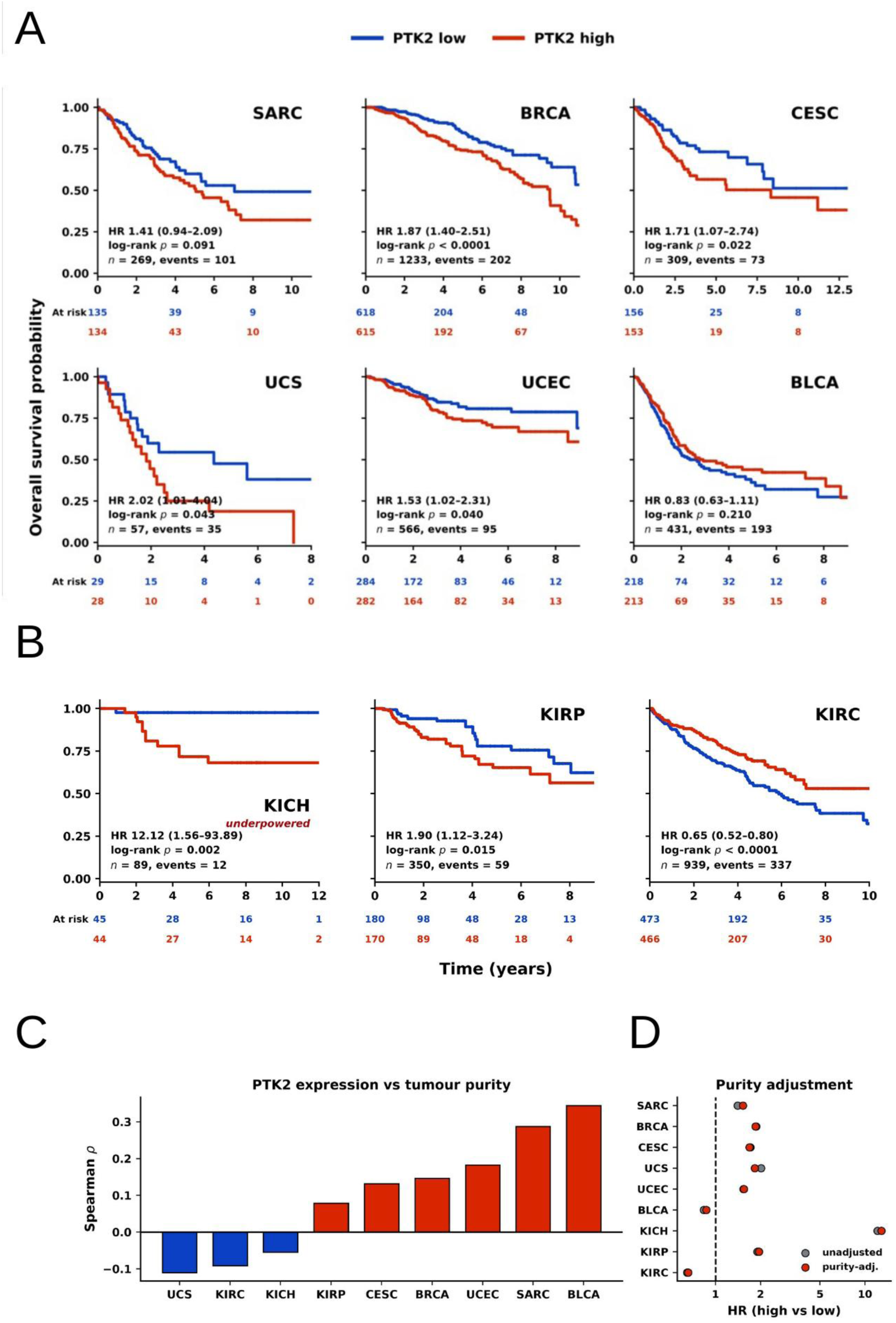
Association of PTK2/FAK mRNA with overall survival and tumour purity. (A) Kaplan-Meier plots for sarcoma, breast, cervical, uterine carcinosarcoma, endometrial and bladder tumours, stratified by PTK2 transcript expression above (red) or below (blue) the cohort median. (B) Kaplan-Meier plots of three renal malignancies: kidney chromophobe (KICH), papillary (KIRP) and clear cell (KIRC) carcinoma. Hazard ratio, 95% confidence interval, log-rank p value, cohort size and event count are displayed in each plot, with numbers at risk below. (C) Spearman correlation between PTK2 expression and ABSOLUTE consensus tumour purity by cohort. (D) Hazard ratios before (grey) and after (red) adjustment for tumour purity. *Alt text: Kaplan-Meier survival curves stratified by PTK2 expression, with scatter plots of PTK2 expression against tumour purity and hazard ratios before and after purity adjustment*.

The renal tumours make the discordance explicit (Figure 5B). Across three subtypes, FAK transcripts were associated with better outcome in kidney renal clear cell carcinoma (KIRC; HR 0.65, 0.52-0.80, p < 0.0001) but poorer outcome in kidney renal papillary cell carcinoma (KIRP; HR 1.90, 1.12-3.24, p = 0.015) and kidney chromophobe cancer (KICH; HR 12.12, 1.56-93.89, p = 0.002). The KICH estimate rests on twelve events in eighty-nine patients and its confidence interval spans nearly two orders of magnitude; it is reported for completeness but is underpowered and not interpreted further. The same gene therefore reverses direction between tumours of the same organ.

Because FAK is expressed by stromal and immune as well as malignant cells, we asked whether these associations reflect tumour purity. The correlation between PTK2 expression and ABSOLUTE consensus purity was weak in every cohort, from -0.11 in uterine carcinosarcoma to +0.34 in bladder carcinoma (Figure 5C), and repeating the survival models with purity as a covariate left hazard ratios essentially unchanged in all nine cohorts, including the reversed association in KIRC (Figure 5D). The reversal is therefore a property of the tumour types themselves, not a proxy for malignant cell content.

### 3.6. A threshold-free re-derivation separates cis from trans PTK2-associated transcripts

The preceding sections leave a specific problem. PTK2 is gained rather than mutated, its transcript reflects that gain and in turn predicts protein, and yet its association with survival runs in opposite directions in different cohorts. Neither coding variation nor tumour purity accounts for the discordance. What remains is the possibility that the transcriptional programme accompanying PTK2 is not itself the same across cancers.

Transcriptomic studies have identified genes expressed in a FAK-dependent manner [43,44], yet the resulting lists are not concordant across cancers, and it has been unclear whether this reflects model systems or tumour types. Many such studies also lack a FAK knockout, so it is not evident whether genes are directly influenced by FAK or are co-expressed with it as part of a wider cellular programme. A further difficulty is that public co-expression summaries report only correlations above a fixed coefficient, so a transcript below threshold in one cohort cannot be distinguished from one never measured.

We therefore recomputed Spearman correlations between PTK2 and every transcript directly from the expression matrix of each of 19 cohorts, retaining all coefficients (Table S5). A transcript entered the consensus where it reached rho at least 0.30 at FDR below 0.05 in at least six of the seventeen cohorts with at least 100 patients, each cohort capped at its 500 strongest correlations per class (Table S2). This yielded 217 transcripts: 173 on chromosome 8q, designated cis, and 44 elsewhere, designated trans (Table S3).

The cis transcripts correlated with PTK2 across nearly all cohorts, forming a dense and largely uninterrupted block (Figure 6A; Figure S3), consistent with regional co-regulation across accessible 8q chromatin. The trans transcripts showed a different structure (Figure 6B): strongly correlated in some cohorts and weakly or not at all in others, with cohorts separating visibly into bands. This separation, rather than the gene set itself, is the principal observation.

**Figure 6.**
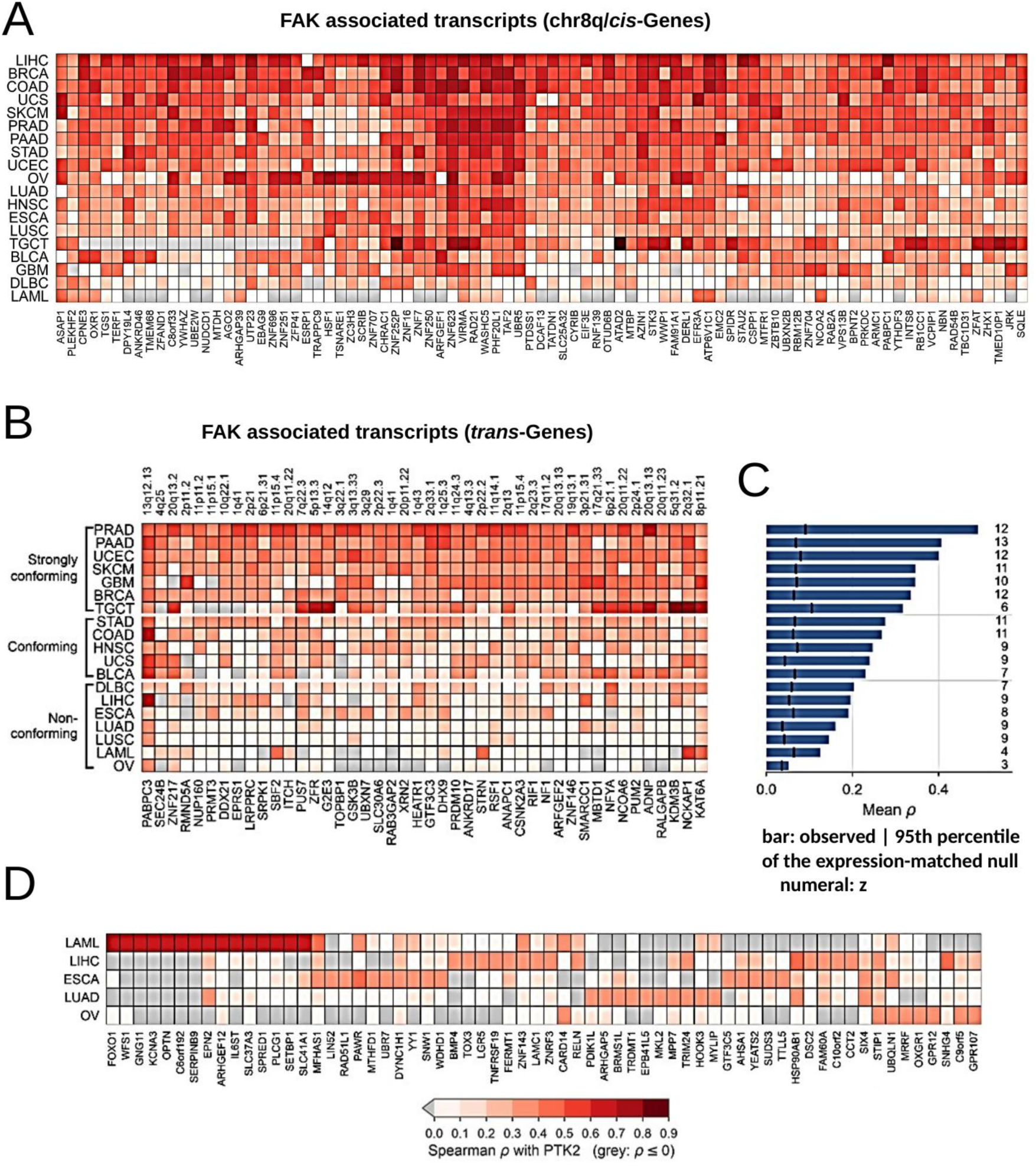
Pan-cancer analysis of transcripts associated with PTK2/FAK. (A) Transcripts on chromosome 8q (cis genes) positively associated with PTK2/FAK expression, plotted as a heat map of Spearman correlations by cancer type; the remaining cis transcripts are shown in Supplementary Figure S3 on the same scale. (B) Transcripts present on chromosomal arms other than 8q (trans) that display an association with PTK2/FAK expression, with cohorts grouped into strongly conforming, conforming and nonconforming tiers. Grey indicates rho <= 0. (C) Mean Spearman correlation between PTK2 and the trans set by cohort (conforming score); the dashed line marks the 95th percentile of the expression-matched null and the numeral gives the z-score. (D) Among the nonconforming tumours, the most highly correlated PTK2-associated transcripts for each cohort, drawn exclusively from transcripts that did not meet the recurrence requirement. *Alt text: Heat maps of Spearman correlation between PTK2 and its cis and trans associated transcripts across cancer cohorts*.

### 3.7. Tumour types stratify into conforming and nonconforming groups

To quantify the separation, each cohort was scored by the mean correlation between PTK2 and the trans set and compared against 1,000 expression-decile-matched random sets (Figure 6C; Table S4). Scores ranged from a mean rho near 0.5 to below 0.1. Strongly conforming cohorts, comprising prostate, pancreatic, uterine corpus endometrial, melanoma, glioblastoma, breast and testicular germ cell tumours, correlated strongly with one another in FAK-associated transcription. Conforming cohorts, comprising stomach, colon, head and neck, uterine carcinosarcoma and bladder, were similar in pattern but more modest. Nonconforming cohorts, including lung, liver, oesophagus, blood and ovary, displayed patterns distinct from the other groups and from each other.

Nonconforming cohorts correlated with the others for cis genes but lacked coordinated trans co-expression. Their absence of the consensus programme does not mean PTK2 lacks correlates there: taking each nonconforming cohort’s most strongly PTK2-correlated transcripts drawn exclusively from genes failing the recurrence requirement shows abundant and strong correlation (Figure 6D; Table S14), but these transcripts are largely private to individual cohorts. PTK2 co-expression in nonconforming tumours is therefore not weaker but different, drawing on partners not shared across tumour types.

### 3.8. The trans programme is not explained by copy number

Because PTK2 lies on a frequently gained arm, co-amplification is an alternative explanation for co-expression, applying with particular force to the cis transcripts. We computed partial Spearman correlations after residualising both PTK2 and each transcript on 8q arm copy number and genome-wide aneuploidy score (Table S6). The cis transcripts serve as the positive control, since they lie on the same arm and must attenuate if the covariate is effective. They did, falling by 38% at the median with attenuation evident in every cohort (Figure S4A). The trans transcripts did not: the median change was +2%, cohort-level scores were essentially unmoved, and strongly conforming and nonconforming cohorts retained their assignments (Figure S4B).

The two classes therefore behave as different phenomena. The cis association is substantially a consequence of regional copy number, as its genomic position predicts, whereas the trans programme is independent of PTK2 dosage and of overall aneuploidy. The conforming and nonconforming distinction is accordingly not a distinction between more and less amplified tumour types.

### 3.9. Co-expression is detectable within individual tumours

Bulk expression mixes malignant, stromal and immune compartments, so a correlation computed across patients can arise from variation in tissue composition rather than from co- expression within cells. We therefore computed co-expression across metacells within each tumour, so that each patient serves as their own control, using single-cell data from tumour types matched to the bulk cohorts. Of 392 tumours with at least 300 malignant cells, 285 had PTK2 detected above the floor, and 243 of these carried at least 20 of the 44 trans transcripts above the floor and were scored.

Of these, 133 (55%) exceeded their own expression-matched null (median z = +1.85; Wilcoxon p = 3.4 x 10-31) (Figure 7A; Table S16). The effect was present across tumour types rather than carried by one (Figure 7B), although unevenly: pancreatic adenocarcinoma contributes 96 tumours and prostate a further 50, whereas cervical (n = 2), ovarian (n = 4) and bladder (n = 7) are too sparsely represented to interpret per-type. Within each tumour the trans transcripts were more strongly correlated with PTK2 than 300 randomly chosen genes (median r 0.081 versus 0.183; paired Wilcoxon p = 4.1 x 10-29) (Figure 7C), confirming specificity to the consensus set rather than to any set of comparable size.

**Figure 7.**
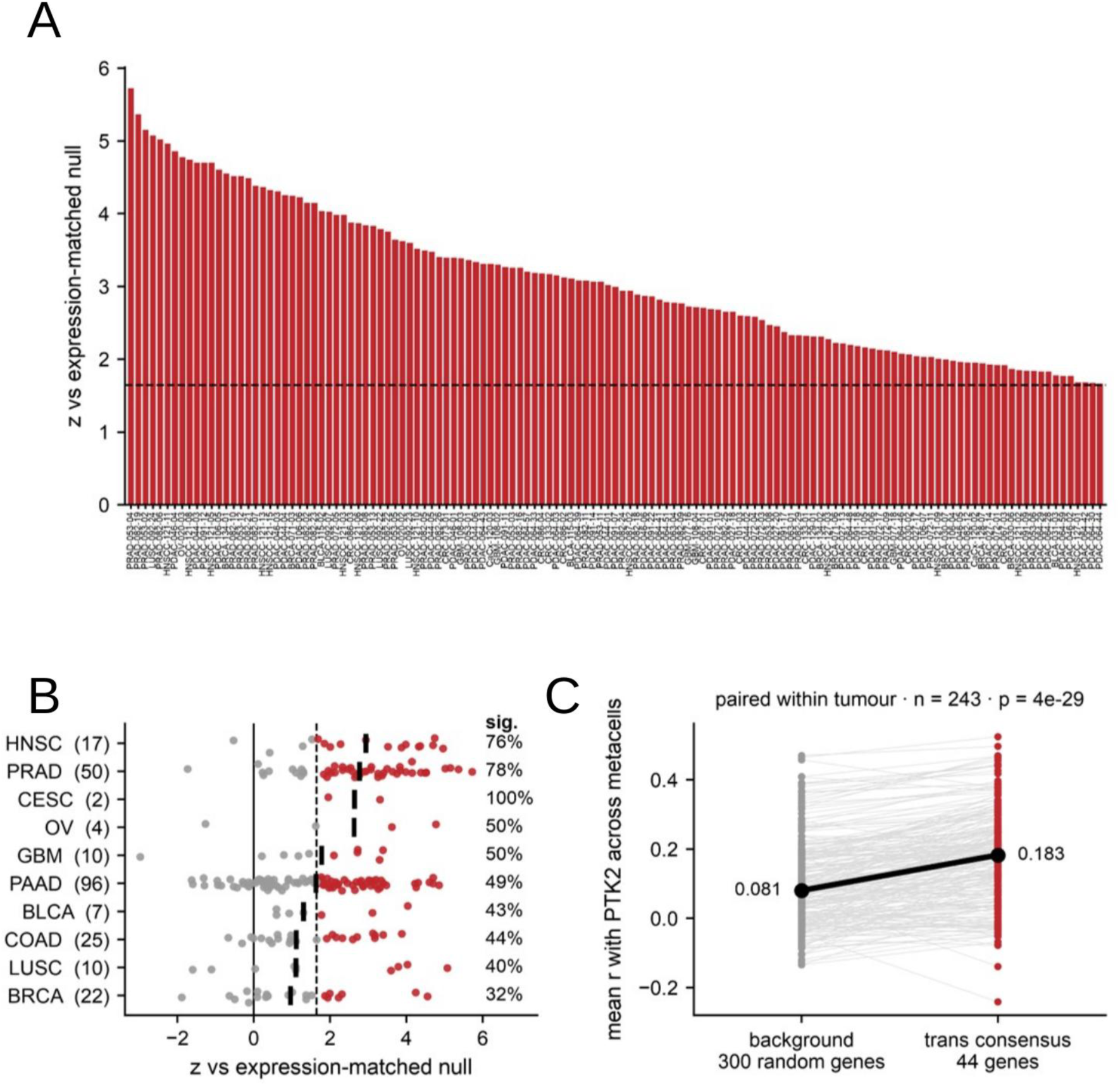
PTK2 co-expression is detectable within individual tumours. (A) Co-expression z against an expression- matched null, computed across metacells within each tumour so that each patient serves as their own control. The 133 tumours reaching p < 0.05 are shown, ordered by z; the dashed line marks the one-sided 0.05 threshold. The 110 tumours not reaching significance are omitted for legibility and are listed with their full statistics in Table S16. (B) The same scores by tumour type, with every scored tumour shown (red, p < 0.05; grey, not significant), the black bar marking the median and the percentage at right giving the fraction significant; numbers in parentheses are tumours per type. (C) Mean correlation with PTK2 across metacells for the 44 trans transcripts compared with 300 randomly chosen genes, paired within each tumour; the black line and markers give the medians. *Alt text: Per-tumour co-expression z scores against expression-matched nulls, a per-cancer-type summary, and a comparison of target against random gene sets*.

Sequencing depth was not limiting: z was unrelated to the number of metacells per tumour (rho = +0.011, p = 0.86) (Figure S5C). The association with PTK2 detection rate was negative (rho = -0.29, p = 3.3 x 10-6) and that with its coefficient of variation across metacells positive (rho = +0.17, p = 0.009) (Figure S5A,B). These describe a measurement constraint rather than a biological one: where PTK2 is detected in nearly all cells it varies little between metacells, leaving little variation against which co-expression can be measured, so 55% is a conservative estimate.

The per-tumour-type ranking does not track the bulk conforming score (rho = -0.07, p = 0.87 across the nine shared cohorts) (Figure S5D). This is expected rather than contradictory: the bulk analysis correlates expression across patients within a cohort, this analysis across cell states within one tumour, and the two capture different sources of variation. The single-cell data establish that PTK2 co-expression is detectable within individual tumours and is not an artefact of bulk composition; they are not intended to reproduce the stratification.

### 3.10. Conforming and nonconforming tumours differ in biology, not in degree

The trans transcripts are distributed non-randomly across the genome. Tested against the universe of measured transcripts, which controls for gene density, they were enriched on 20q (6 observed against 0.8 expected, q = 0.002), 11p (4 against 0.8, q = 0.020) and 2p (5 against 1.1, q = 0.015), with 3q, 2q and 1q not reaching significance (Figure 8A; Table S7). The counts are small and are reported as observed against expected rather than as fold enrichment alone. Pathway analysis used preranked gene set enrichment within each cohort over all measured transcripts, imposing no threshold, and the tiers were then contrasted (Tables S8, S9). The groups were not separated by the degree to which one set of pathways was expressed; they were enriched for different biology, and in one case in opposite directions (Figure 8B). Strongly conforming cohorts were enriched for DNA replication fidelity, histone H3-K4 methylation, replication fork processing, DNA damage checkpoint signalling, histone lysine methylation, histone H4 acetylation, DNA recombination, recombinational repair and interstrand cross-link repair, with recombinational repair and replication fork processing significant in all seven.

**Figure 8.**
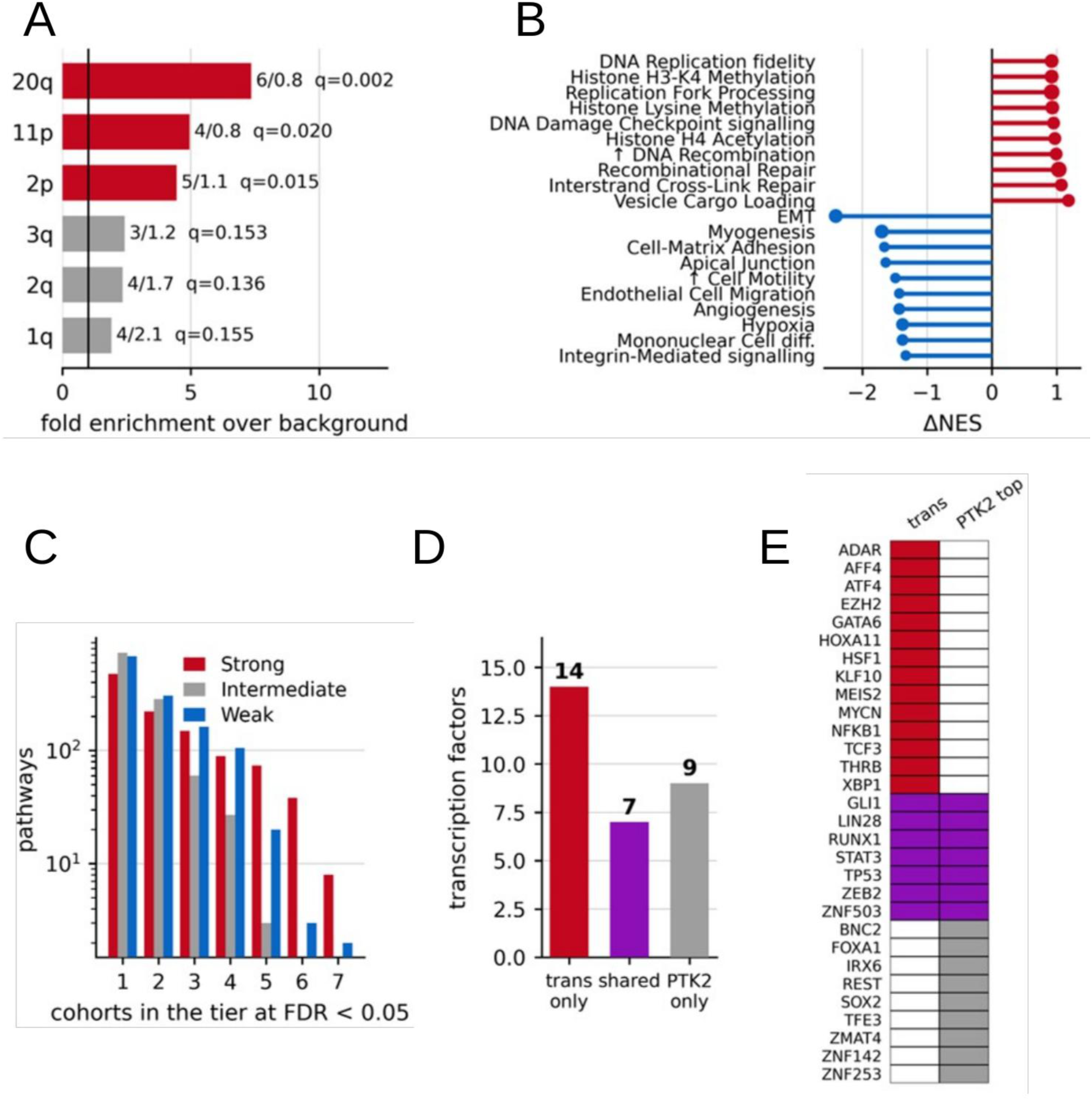
Conforming and nonconforming tumours differ in associated biology. (A) Distribution of the trans transcripts across chromosome arms, tested against the universe of transcripts measured rather than the whole genome. Bars give fold enrichment; labels give observed against expected counts and the Benjamini-Hochberg q value. Red, q < 0.05. (B) Difference in mean normalised enrichment score (NES) between strongly conforming and nonconforming cohorts for the most strongly separating pathways. Red, higher in conforming tumours; blue, higher in nonconforming tumours. (C) Number of pathways reaching FDR < 0.05 in k cohorts of each tier; note the logarithmic axis. (D) Transcription factors enriched in the trans programme only, in both sets, or in PTK2’s top correlates only. (E) Membership of each factor; a filled cell indicates enrichment in that set. *Alt text: Chromosome-arm enrichment bars, pathway enrichment scores contrasting conforming and nonconforming cohorts, and transcription factor enrichment results*.

Nonconforming cohorts were enriched instead for the canonical adhesion biology of FAK: epithelial-to-mesenchymal transition, myogenesis, cell-matrix adhesion, apical junction, cell motility, endothelial cell migration, angiogenesis, hypoxia and integrin-mediated signalling (Figure 8B). Epithelial-to-mesenchymal transition reversed in sign between the groups, with a normalised enrichment score of -1.37 in strongly conforming against +1.04 in nonconforming cohorts, significant in five of seven. The tumour types in which the recurrent PTK2 programme is most coherent are therefore not those in which FAK’s best-characterised functions are most prominent.

The intermediate tier is not a third biological group. Counting pathways reaching FDR below 0.05 in k cohorts of a tier, strongly conforming cohorts share 46 pathways across six or more of their seven members, nonconforming cohorts share five, and the intermediate tier shares none at any comparable recurrence (Figure 8C; Table S10). These cohorts occupy a position on a gradient rather than forming a class.

Finally, we asked whether the trans programme carries a distinct regulatory input relative to PTK2’s general correlates. Transcription factor enrichment was computed for the trans consensus and, separately, for PTK2’s strongest pan-cancer correlates matched in size and built without the recurrence requirement (Tables S11, S12). Of the factors enriched, 14 were specific to the trans programme, 9 to PTK2’s top correlates, and 7 shared (Figure 8D,E). The trans programme therefore carries associations, including ATF4, EZH2, HSF1, MYCN, NFKB1 and XBP1, not recovered from PTK2’s general correlates, consistent with expression within a broader coordinated programme rather than as a consequence of PTK2 abundance.

## 4. Discussion

The practical consequence concerns how conflicting reports about a gene should be read. PTK2 is an adverse prognostic marker in some cohorts and a favourable one in others, and that reversal is not an artefact of tumour purity. Under the conventional reading it is a problem to be explained away by cohort size, platform or chance. The analysis here supports a different reading: the cohorts in which PTK2 behaves differently are also the cohorts in which it keeps different transcriptional company, and the two tiers are enriched for distinct biology rather than for more or less of the same. A gene is therefore not well described as good or bad across cancers, and a meta-analysis that pools cohorts to settle the question averages over the very structure that explains it. Where a gene shows discordant associations, testing whether its co- expression programme is conserved is a more informative response than increasing n.

FAK expression has been correlated with differing suites of transcripts and proteins in different cancers [33–48], and it has been unclear whether this reflects particular model systems or genuine differences between cancers. The tiering offers an answer for one such case: the very different profiles of co-expressed proteins reported in ovarian and pancreatic carcinoma correspond to cohorts on opposite sides of the split, pancreatic adenocarcinoma being strongly conforming and ovarian carcinoma nonconforming.

A methodological point underlies this. Co-expression summaries that report only coefficients above a fixed threshold censor the data rather than summarising it, and a consensus assembled from them cannot distinguish a weak correlation from a missing measurement. Recomputing from the expression matrices changes which transcripts enter the consensus, and symbol resolution alone recovered 66 chromosome 8q transcripts that would otherwise have been misassigned as distal.

The separation of cis from trans transcripts is the second point. PTK2’s position on a frequently gained arm has long complicated the interpretation of FAK expression, and the partial correlation quantifies that complication: approximately 38% of the cis association is attributable to copy number, whereas the trans programme is independent of it. The cis transcripts are not uninteresting, but they are substantially a consequence of genomic position. It is the use of 8q transcripts as a positive control rather than as a result that makes the trans finding interpretable; an unattenuated trans signal would carry little weight without the accompanying demonstration that the covariate is effective.

The third and least expected point concerns the distinction itself. If the recurrent programme marked FAK’s canonical activity, the cohorts expressing it most coherently should be those in which adhesion, motility and epithelial-to-mesenchymal transition dominate. The opposite holds. Conforming cohorts are enriched for recombinational repair, replication fork processing and chromatin modification, whereas adhesion and epithelial-to-mesenchymal transition characterise the cohorts in which the programme is absent, the latter reversing in sign between the groups. One interpretation is that the recurrent programme reflects FAK’s nuclear and transcriptional functions rather than its adhesion-associated signalling, consistent with its independence from copy number. An alternative is that where adhesion signalling dominates, FAK’s partners are set by local tissue context and so do not recur across tumour types, which is what the private correlates of the nonconforming cohorts suggest. Direct support for the adhesion reading comes from ovarian carcinoma, nonconforming here, in which tumour FAK sustains fibrillar collagen expression in ascites tumour cells and FAK inhibition lowers COL1A1, COL3A1 and COL5A2 [9]. These readings are not mutually exclusive and the present data do not distinguish them.

FAK may not be essential for the expression of transcripts with which it is associated. Among twelve rare tumours bearing PTK2 double deletions we identified no consistently lost transcripts, though a called double deletion need not reflect complete loss given heterogeneity and polyploidy. Only reconstitution studies can establish where FAK drives a pathway [47]. FAK is often linked to malignancy, yet PTK2 shows no enrichment for mutations that enhance catalytic activity; instead the gene is frequently gained and transcript and protein are more abundant. This may reflect the importance of noncatalytic scaffolding function, though that seems unlikely given the lack of tumour support from kinase-dead FAK [42] and the activity of kinase inhibitors in vivo [1]. Cycling between active and inactive states may matter more, consistent with a biosensory role rather than a simple proliferative driver [1].

FAK gain is linked to elevated protein here and elsewhere [45], with amplification raising FAK mRNA in colon, breast, prostate and hepatocellular carcinoma [46] and gene dosage tracking protein in head and neck squamous carcinoma; late-stage ovarian cells show higher FAK protein and kinase-reporter activity [47]. Protein is elevated in tumour relative to normal tissue across many cancers [52–55]. In the present analysis FAK was an unfavourable prognostic marker in breast, cervical, kidney renal papillary, uterine carcinosarcoma and uterine corpus endometrial carcinoma, with sarcoma trending in the same direction without reaching significance.

These results align with reports that elevated FAK expression and activation predict decreased survival [48], particularly in ovarian and breast cancer [1,45]. The opposing prognostic direction in renal cancer may stem from differences in microenvironment, molecular subtype and oncogenic dependency: KIRC is driven principally by VHL mutation [11,12,49] and hypoxia-related pathways [50], and may therefore not depend on FAK to sustain those signals. The purity-adjusted analysis indicates these opposing associations are not explained by differences in malignant cell content. In bladder cancer we found no significant association with overall survival, in contrast to prior reports linking elevated PTK2 to advanced stage and shorter survival [51].

Transcription factor analysis identified regulatory associations specific to the recurrent programme, including EZH2, which promotes progression through gene silencing [56], alongside ATF4, HSF1, MYCN, NFKB1 and XBP1. This is consistent with the recurrent programme being expressed within a broader cellular programme rather than as a direct readout of FAK abundance. Among factors associated with PTK2’s general correlates rather than the recurrent programme was REST, which exhibits dual roles in tumour growth and suppression [57–59].

The framework has its own scope conditions. It assumes co-expression computed within a cohort is meaningful, which requires cohorts large enough for stable rank correlation; below roughly one hundred patients the permutation null dominates the variance and tier assignment is unreliable. It assumes copy number is measured on the same patients as expression, which holds for TCGA but not every resource. The cis control is informative only for genes in regions of recurrent copy-number change; for a gene in a stable region there is nothing for the adjustment to remove, and its failure to remove anything must not be reported as evidence of dosage-independence. Finally, the tiers are properties of the index gene together with the cohort set examined, not labels that transfer between genes.

Several limitations apply. The analysis is observational throughout: co-expression constrains which transcripts accompany PTK2 but does not establish direction, and causal attribution would require perturbation that public data do not support. The single-cell analysis addresses compartment mixing but rests on 243 tumours from ten tumour types, with pancreatic adenocarcinoma and prostate together contributing 60% of those scored. A further 42 tumours could not be scored because fewer than 20 of the 44 transcripts cleared the detection floor, and where PTK2 is detected in nearly all cells its expression varies too little between metacells for co-expression to be measurable, so the proportion reported is conservative. The copy-number control uses arm-level calls and a genome-wide aneuploidy score rather than focal gene-level copy number, the appropriate resolution for an arm-level event but not for a focal alteration at a distal locus. The tiers are defined on a continuous score, and while the extremes are well separated and biologically distinct, the intermediate tier is a position on that gradient rather than a discrete class.

The first FAK-targeting drugs have entered the clinic and more are likely to follow. Combination strategies are emerging: in ovarian cancer, FAK inhibition with pegylated doxorubicin and anti-TIGIT enhances survival and tertiary lymphoid structure formation, traced to release of omega-3 fatty acids and consequent CXCL13 production by resident peritoneal macrophages [8]. Since these drugs exhibit tumour-specific effects, differences in tumour FAK biology of the kind described here will matter. Dissecting the signalling webs associated with PTK2 across and within tumours may provide contextual insight for the use of FAK inhibitors and identify biomarker candidates reporting FAK-associated programmes in patients.

## 5. Conclusions

PTK2/FAK is rarely mutated but frequently gained through arm-level events on chromosome 8q, and this gain is transcriptionally and translationally consequential across most solid tumours. Re-deriving PTK2 co-expression without a correlation threshold identifies 217 recurrent transcripts that partition cleanly by genomic position: a 173-transcript cis set on 8q that is substantially attributable to copy number, and a 44-transcript trans set that is not. Tumour types differ markedly in how coherently they express the trans programme, and the difference is one of biology rather than magnitude, with conforming tumours enriched for replication and repair processes and nonconforming tumours for the adhesion and epithelial-to-mesenchymal programmes with which FAK is classically associated. This distinction is a candidate axis along which the molecular context for FAK-directed therapy might be defined.

## Supplementary data

Supplementary data are available at Briefings in Bioinformatics online. Supplementary Tables S1 to S16 are provided as a single Microsoft Excel workbook (Supplementary_Tables.xlsx) containing one worksheet per table, each worksheet named for its table number (S1_Sources, S2_Parameters, and so on), together with a Contents worksheet listing every table with its legend. Supplementary Figure S1: chromosome 8 copy number in liver, colon and stomach adenocarcinoma; Supplementary Figure S2: PTK2 expression across tissue state and pathological stage; Supplementary Figure S3: additional chromosome 8q (cis) transcripts; Supplementary Figure S4: co-expression is not explained by copy number; Supplementary Figure S5: single-cell diagnostics and comparison with the bulk analysis; Table S1: data sources and software; Table S2: gene-set definition parameters; Table S3: the frozen PTK2 co- expression gene set; Table S4: per-cohort co-expression scores with expression-matched nulls; Table S5: PTK2 correlation matrix for the frozen gene set; Table S6: partial correlation controlling for copy number; Table S7: chromosome-arm distribution of the trans programme; Table S8: pathways separating conforming from nonconforming cohorts; Table S9: preranked GSEA results per cohort; Table S10: pathway sharing within each tier; Table S11: transcription-factor enrichment; Table S12: transcription factors by set membership; Table S13: retired to current gene symbols; Table S14: transcripts of the nonconforming cohorts; Table S15: overall survival by PTK2 expression; Table S16: per-tumour single-cell co- expression results.

## Funding

This work was supported by the National Institutes of Health [R01CA247562 to D.G.S.]; and by a pilot award from the Cancer Cell Map Initiative [U54CA274502].

## Conflict of interest

None declared.

## Author contributions

Conceptualization, S.U.K. and D.G.S.; methodology, S.U.K.; software, S.U.K.; validation, S.U.K.; formal analysis, S.U.K.; investigation, S.U.K.; data curation, S.U.K.; writing — original draft preparation, S.U.K.; writing — review and editing, S.U.K., A.S., A.M.T. and D.G.S.; visualization, S.U.K.; supervision, D.G.S. and A.M.T.; project administration, D.G.S.; funding acquisition, D.G.S. and A.M.T. All authors have read and agreed to the submitted version of the manuscript.

## Data availability

All primary data are publicly available from the sources listed in Table S1. The single-cell datasets analysed here are available under the following accessions. Gene Expression Omnibus: GSE135337, GSE143791, GSE173468, GSE181294, GSE185965, GSE188807, GSE193766, GSE201348, GSE202051, GSE208653, GSE210358, GSE211644, GSE212966, GSE213047, GSE217956, GSE222300, GSE224679, GSE225600, GSE226762, GSE231535, GSE232525, GSE234832, GSE242230, GSE243359, GSE244267, GSE249263. ArrayExpress: E-MTAB-8107, E-MTAB-8410, E-MTAB-8559, E-MTAB-12022, E-MTAB- 12528. BioProject: PRJCA001063, PRJCA008495, PRJCA016878, PRJNA879876, PRJNA976462. Derived data, including the frozen gene set, per-cohort scores and the copy- number control, are provided in the supplementary workbook. The final analysis code and the genome-wide correlation matrix will be made available at the GitHub repository of S.U.K. (https://github.com/Safir-Ullah-Khan) upon acceptance of the manuscript.

## Key Points

- Conflicting reports of a gene’s prognostic direction across cancers may reflect genuine lineage-dependent differences in its transcriptional context rather than irreproducibility, and this is an empirical claim that can be tested.
- Public co-expression summaries truncate at a fixed coefficient, so a gene below threshold in one cohort cannot be distinguished from one never measured; recomputing correlations threshold-free from the expression matrices removes this bias.
- A gene in a recurrently gained region can acquire correlates through dosage alone; residualising copy number while using the gene’s own chromosomal neighbours as a positive control shows the adjustment works before the result is interpreted.
- Applied to PTK2/FAK, whose association with survival reverses direction between cohorts, the framework separates tumour types into conforming and nonconforming groups enriched for different biology rather than differing in degree.
- A within-tumour single-cell test recovers co-expression in 133 of 243 tumours and shows that bulk and single-cell analyses interrogate different variance components, which constrains how pan-cancer co-expression results should be read.

## Author descriptions

Safir Ullah Khan is a postdoctoral researcher at Moores Cancer Center, University of California San Diego. He holds a PhD in cell biology and works on computational oncology and pan-cancer transcriptomics.

Ava Storgard is a researcher in the Stupack laboratory at Moores Cancer Center, University of California San Diego. Her main research interest is cancer bioinformatics.

Alfonso Mendez Tenorio is a Full Professor at the National Polytechnic Institute, Mexico City, where he is principal investigator of the Laboratory of Bioinformatics and Biotechnology.

Dwayne G. Stupack is a Full Professor and principal investigator at Moores Cancer Center, University of California San Diego. His research combines laboratory and computational oncology, with a focus on gynaecological cancers.

## Supplementary Figure Legends

**Figure S1.** Chromosome 8 copy number in additional tumour types. CAIRN-based representation of copy number along chromosome 8 in individual tumours of hepatocellular carcinoma (LIHC), colon adenocarcinoma (COAD) and stomach adenocarcinoma (STAD), plotted as in Figure 2B.

**Figure S2.** PTK2 expression across tissue state and pathological stage. (A) PTK2 expression in normal (N), primary tumour (T) and metastatic (M) tissue in breast and head and neck cancer; ****, p < 0.0001. (B) PTK2 expression by pathological stage in six cohorts, with Spearman correlation coefficient, p value and sample number given on each panel.

**Figure S3.** Additional chromosome 8q (cis) transcripts. PTK2 co-expression with the remaining cis transcripts of the frozen consensus, on the same colour scale and cohort ordering as Figure 6A. Together the two panels cover the complete cis set. Grey indicates rho <= 0.

**Figure S4.** Co-expression is not explained by copy number. Mean Spearman correlation between PTK2 and each transcript set by cohort, before (open circles) and after (filled symbols) partial correlation against 8q arm copy number and genome-wide aneuploidy score. (A) Chromosome 8q (cis) transcripts, the positive control, which attenuate by 38% on median. (B) The trans consensus, which does not attenuate (median change +2%). Symbol shape denotes cohort tier.

**Figure S5.** Single-cell diagnostics and comparison with the bulk analysis. (A) Co-expression z against PTK2 detection rate, coloured by the coefficient of variation of PTK2 across metacells. (B) z against that coefficient of variation. (C) z against the number of metacells per tumour. (D) Median single-cell z per tumour type against the corresponding bulk cohort score, for the nine cohorts present in both analyses. Spearman rho and p are given on each panel.

## Notes

https://github.com/Safir-Ullah-Khan/Pan-cancer-framework-for-conforming-and-nonconforming-co-expression-programs

### Competing Interest Statement

The authors have declared no competing interest.

## References

1. Schlaepfer DD, Ojalill M, Stupack DG. Focal adhesion kinase signaling—tumor vulnerabilities and clinical opportunities. J Cell Sci 2024;137:jcs261723.

2. Chen Y, et al. Identification of PTK2 as an adverse prognostic biomarker in breast cancer by integrated bioinformatics and experimental analyses. Front Mol Biosci 2022;9:984564.

3. Liu L-L, et al. Proteomic Comparison of the Cytotoxicology of Two Diastereomers of Citreamicin Reveals Differentially Activation of NF-κB Pathway. Front Mar Sci 2020;7:228.

4. Dawson JC, et al. Targeting FAK in anticancer combination therapies. Nat Rev Cancer 2021;21:313–324.

5. Genna A, et al. FAK family proteins regulate in vivo breast cancer metastasis via distinct mechanisms. bioRxiv 2023;2023.10.27.564212.

6. Hu H-H, et al. Roles and inhibitors of FAK in cancer: current advances and future directions. Front Pharmacol 2024;15:1274209.

7. Tai Y-L, et al. An EGFR/Src-dependent β4 integrin/FAK complex contributes to malignancy of breast cancer. Sci Rep 2015;5:16408.

8. Chen XL, Tharp KM, Ojalill M, Ozmadenci D, Boyer A, Haanen TJ, et al. FAK inhibition in ovarian cancer releases omega-3 fatty acids to program CXCL13-producing anti-tumor resident peritoneal macrophages. Cell Rep 2026;45:117009.

9. Ojalill M, Haanen TJ, Khan SU, Zhang Y, Boyer A, Chen XL, Suwal U, Rappu P, Heino J, McHale MT, et al. Tumor FAK supports fibrillar collagen expression in ovarian cancer ascites. Matrix Biol 2026;149:102053.

10. Shi Y, Zhang Y, Chen Y, et al. Comprehensive pan-cancer analysis of 33 human cancers reveals immunotherapeutic value of focal adhesion tyrosine kinase. Medicine (Baltimore) 2024;103:e37362.

11. Cerami E, et al. The cBio cancer genomics portal: an open platform for exploring multidimensional cancer genomics data. Cancer Discov 2012;2:401–404.

12. Gao J, et al. Integrative analysis of complex cancer genomics and clinical profiles using the cBioPortal. Sci Signal 2013;6:pl1.

13. Bartha Á, Győrffy B. TNMplot.com: a web tool for the comparison of gene expression in normal, tumor and metastatic tissues. Int J Mol Sci 2021;22:2622.

14. Edgar R, Domrachev M, Lash AE. Gene Expression Omnibus: NCBI gene expression and hybridization array data repository. Nucleic Acids Res 2002;30:207–210.

15. Barrett T, et al. NCBI GEO: archive for functional genomics data sets—update. Nucleic Acids Res 2013;41:D991–D995.

16. Ru B, et al. TISIDB: an integrated repository portal for tumor-immune system interactions. Bioinformatics 2019;35:4200–4202.

17. Chandrashekar DS, et al. UALCAN: An update to the integrated cancer data analysis platform. Neoplasia 2022;25:18–27.

18. Uhlén M. A human protein atlas for normal and cancer tissues. In Proceedings of the 10th International Conference on Miniaturized Systems for Chemistry and Life Sciences (MicroTAS 2006), Tokyo, Japan, 5–9 November 2006.

19. Goldman MJ, Craft B, Hastie M, Repečka K, McDade F, Kamath A, Banerjee A, Luo Y, Rogers D, Brooks AN, et al. Visualizing and interpreting cancer genomics data via the Xena platform. Nat Biotechnol 2020;38:675–678.

20. Liu J, Lichtenberg T, Hoadley KA, Poisson LM, Lazar AJ, Cherniack AD, Kovatich AJ, Benz CC, Levine DA, Lee AV, et al. An integrated TCGA pan-cancer clinical data resource to drive high- quality survival outcome analytics. Cell 2018;173:400–416.e11.

21. Carter SL, Cibulskis K, Helman E, McKenna A, Shen H, Zack T, Laird PW, Onofrio RC, Winckler W, Weir BA, et al. Absolute quantification of somatic DNA alterations in human cancer. Nat Biotechnol 2012;30:413–421.

22. Taylor AM, Shih J, Ha G, Gao GF, Zhang X, Berger AC, Schumacher SE, Wang C, Hu H, Liu J, et al. Genomic and functional approaches to understanding cancer aneuploidy. Cancer Cell 2018;33:676–689.e3.

23. Seal RL, Braschi B, Gray K, Jones TEM, Tweedie S, Haim-Vilmovsky L, Bruford EA. Genenames.org: the HGNC resources in 2023. Nucleic Acids Res 2023;51:D1003–D1009.

24. Benjamini Y, Hochberg Y. Controlling the false discovery rate: a practical and powerful approach to multiple testing. J R Stat Soc Series B Stat Methodol 1995;57:289–300.

25. Subramanian A, Tamayo P, Mootha VK, Mukherjee S, Ebert BL, Gillette MA, Paulovich A, Pomeroy SL, Golub TR, Lander ES, et al. Gene set enrichment analysis: a knowledge-based approach for interpreting genome-wide expression profiles. Proc Natl Acad Sci USA 2005;102:15545–15550.

26. Liberzon A, Birger C, Thorvaldsdóttir H, Ghandi M, Mesirov JP, Tamayo P. The Molecular Signatures Database hallmark gene set collection. Cell Syst 2015;1:417–425.

27. Fang Z, Liu X, Peltz G. GSEApy: a comprehensive package for performing gene set enrichment analysis in Python. Bioinformatics 2023;39:btac757.

28. Chen EY, et al. Enrichr: interactive and collaborative HTML5 gene list enrichment analysis tool. BMC Bioinform 2013;14:1–14.

29. Xie Z, et al. Gene set knowledge discovery with Enrichr. Curr Protoc 2021;1:e90.

30. Kuleshov MV, et al. Enrichr: a comprehensive gene set enrichment analysis web server 2016 update. Nucleic Acids Res 2016;44:W90–W97.

31. Vogelstein B, et al. Cancer genome landscapes. Science 2013;339:1546–1558.

32. Forbes SA, et al. The Catalogue of Somatic Mutations in Cancer (COSMIC). Curr Protoc Hum Genet 2008;Chapter 10:Unit 10.11.

33. Jacamo RO, Rozengurt E. A truncated FAK lacking the FERM domain displays high catalytic activity but retains responsiveness to adhesion-mediated signals. Biochem Biophys Res Commun 2005;334:1299–1304.

34. Cooper LA, Shen TL, Guan JL. Regulation of focal adhesion kinase by its amino-terminal domain through an autoinhibitory interaction. Mol Cell Biol 2003;23:8030–8041.

35. Cohen LA, Guan JL. Residues within the first subdomain of the FERM-like domain in focal adhesion kinase are important in its regulation. J Biol Chem 2005;280:8197–8207.

36. Cai X, et al. Spatial and temporal regulation of focal adhesion kinase activity in living cells. Mol Cell Biol 2008;28:201–214.

37. Goni GM, et al. Phosphatidylinositol 4,5-bisphosphate triggers activation of focal adhesion kinase by inducing clustering and conformational changes. Proc Natl Acad Sci USA 2014;111:E3177–E3186.

38. Choi CH, et al. pH sensing by FAK-His58 regulates focal adhesion remodeling. J Cell Biol 2013;202:849–859.

39. Acebron I, et al. Structural basis of Focal Adhesion Kinase activation on lipid membranes. EMBO J 2020;39:e104743.

40. Le Coq J, et al. New insights into FAK structure and function in focal adhesions. J Cell Sci 2022;135:jcs259089.

41. Delaney JR, et al. Autophagy gene haploinsufficiency drives chromosome instability, increases migration, and promotes early ovarian tumors. PLoS Genet 2020;16:e1008558.

42. Diaz Osterman CJ, et al. FAK activity sustains intrinsic and acquired ovarian cancer resistance to platinum chemotherapy. eLife 2019;8:e47327.

43. Chuang H-H, et al. FAK in cancer: from mechanisms to therapeutic strategies. Int J Mol Sci 2022;23:1726.

44. Dimitrakopoulos C, et al. Network-based integration of multi-omics data for prioritizing cancer genes. Bioinformatics 2018;34:2441–2448.

45. Sulzmaier FJ, Jean C, Schlaepfer DD. FAK in cancer: mechanistic findings and clinical applications. Nat Rev Cancer 2014;14:598–610.

46. Canel M, et al. Overexpression of focal adhesion kinase in head and neck squamous cell carcinoma is independent of fak gene copy number. Clin Cancer Res 2006;12:3272–3279.

47. Osterman CJD, et al. Abstract A61: FAK activity sustains intrinsic and acquired ovarian cancer resistance to platinum chemotherapy. Clin Cancer Res 2020;26:A61.

48. Zeng X-Q, et al. Prognostic value of focal adhesion kinase (FAK) in human solid carcinomas: a meta-analysis. PLoS ONE 2016;11:e0162666.

49. De Bruijn I, et al. Analysis and visualization of longitudinal genomic and clinical data from the AACR project GENIE biopharma collaborative in cBioPortal. Cancer Res 2023;83:3861–3867.

50. Liu T, et al. Hypoxia-induced PLOD2 promotes clear cell renal cell carcinoma progression via modulating EGFR-dependent AKT pathway activation. Cell Death Dis 2023;14:774.

51. Kim C, et al. Unveiling Bladder Cancer Prognostic Insights by Integrating Patient-Matched Sample and CpG Methylation Analysis. Medicina 2024;60:1175.

52. Miyazaki T, et al. FAK overexpression is correlated with tumour invasiveness and lymph node metastasis in oesophageal squamous cell carcinoma. Br J Cancer 2003;89:140–145.

53. Zhang Z, et al. Functional and clinical characteristics of focal adhesion kinases in cancer progression. Front Cell Dev Biol 2022;10:1040311.

54. Murphy JM, et al. Targeting focal adhesion kinase in cancer cells and the tumor microenvironment. Exp Mol Med 2020;52:877–886.

55. Brown NF, et al. A study of the focal adhesion kinase inhibitor GSK2256098 in patients with recurrent glioblastoma with evaluation of tumor penetration of [11C] GSK2256098. Neuro Oncol 2018;20:1634–1642.

56. Chang C, Hung M. The role of EZH2 in tumour progression. Br J Cancer 2012;106:243–247.

57. Negrini S, et al. REST: an oncogene or a tumor suppressor?. Trends Cell Biol 2013;23:289–295.

58. Singh A, et al. REST-dependent downregulation of von Hippel-Lindau tumor suppressor promotes autophagy in SHH-medulloblastoma. Sci Rep 2024;14:13596.

59. Huang Z, Bao S. Ubiquitination and deubiquitination of REST and its roles in cancers. FEBS Lett 2012;586:1602–1605.

