## Supplementary tables and Fig's for "Context, not contradiction: a pan-cancer framework for conforming and nonconforming co-expression programmes": Supplementary_Data_BriefBioinform.pdf

### Contents

Supplementary Figures S1 to S5 are reproduced below with their legends.

Supplementary Tables S1 to S16 are provided as a separate Microsoft Excel workbook (Supplementary_Tables.xlsx), one worksheet per table, with a Contents worksheet carrying every legend. The legends are also listed at the end of this document.

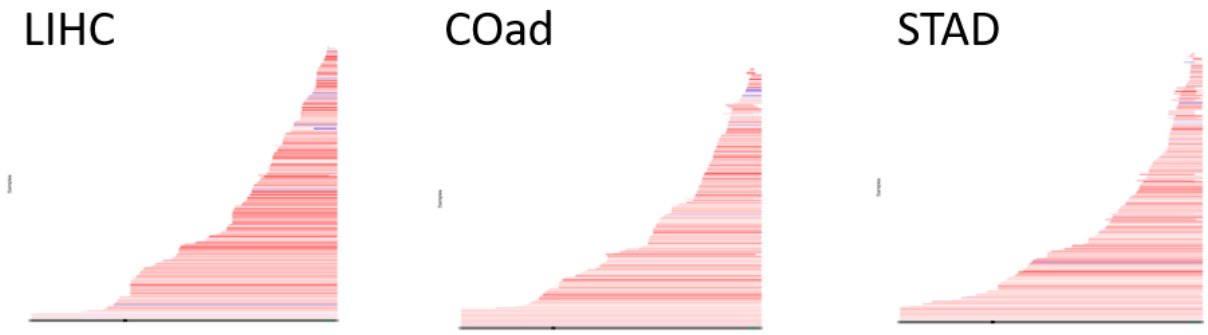


**Figure S1. Chromosome 8 copy number in additional tumour types.** CAIRN-based representation of copy number along chromosome 8 in individual tumours of hepatocellular carcinoma (LIHC), colon adenocarcinoma (COAD) and stomach adenocarcinoma (STAD), plotted as in Figure 2B. Each horizontal line is one tumour, ordered by extent of gain. Red indicates gain and blue loss.

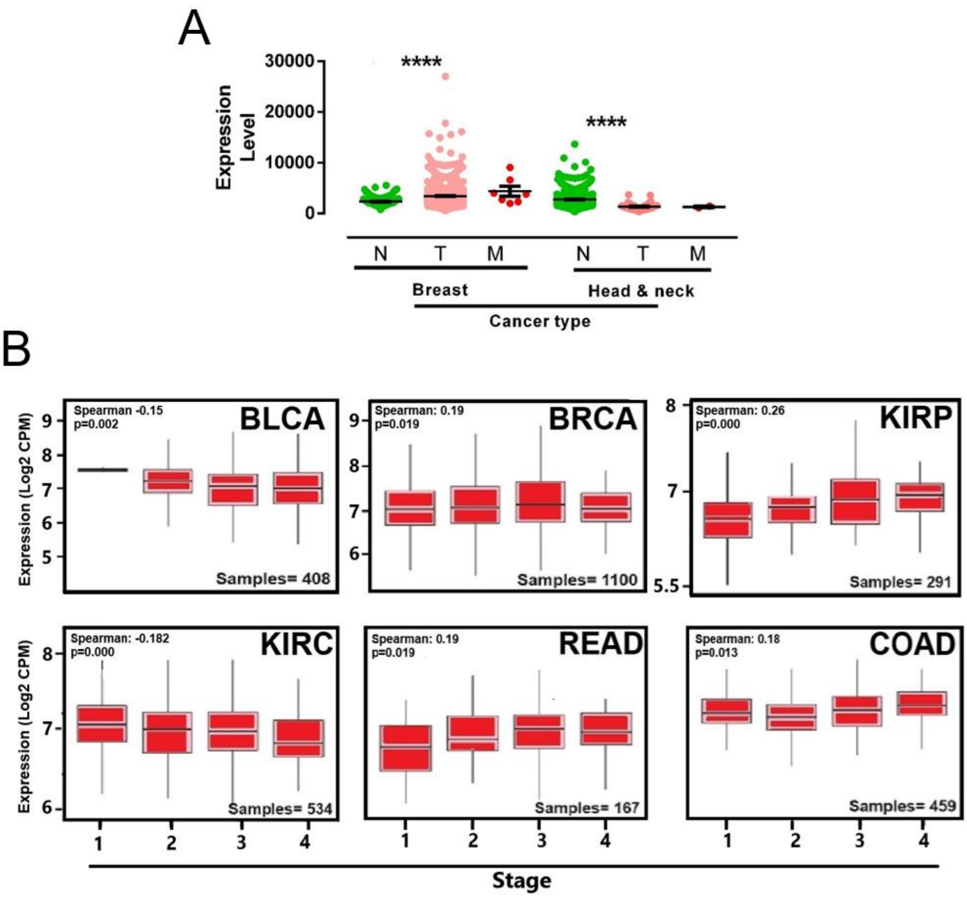


**Figure S2. PTK2 expression across tissue state and pathological stage.** (A) PTK2 expression in normal (N), primary tumour (T) and metastatic (M) tissue in breast and head and neck cancer; ****, $p < 0.0001$. (B) PTK2 expression by pathological stage in six cohorts, with the Spearman correlation coefficient, p value and sample number given on each panel. No consistent trend with stage is evident across cohorts.

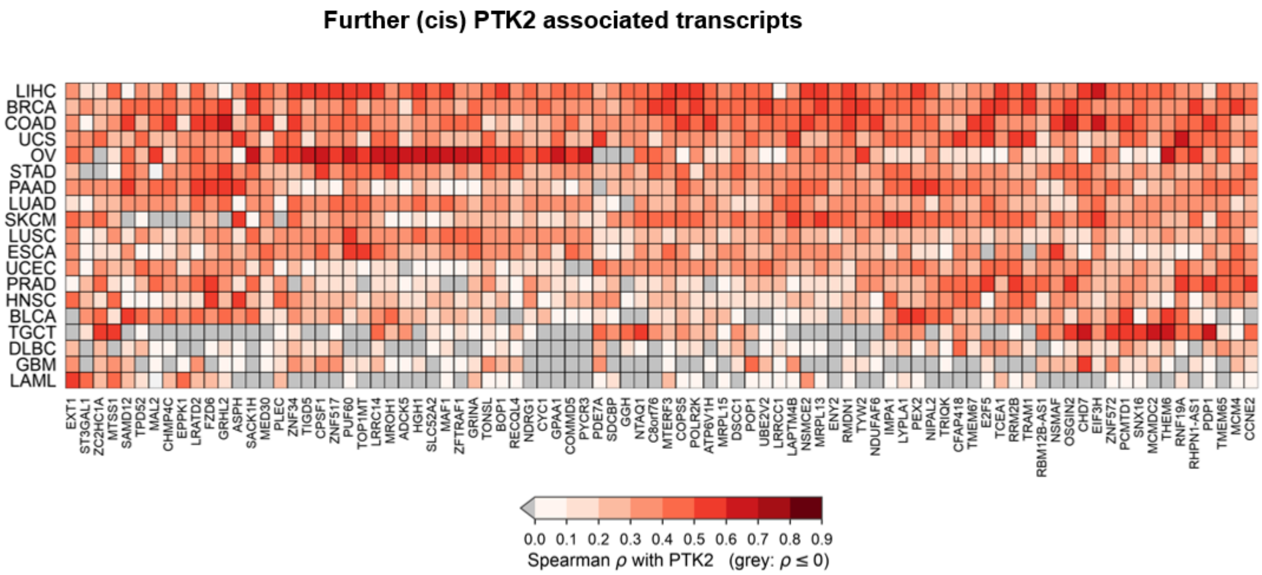


**Figure S3. Additional chromosome 8q (cis) transcripts.** PTK2 co-expression with the remaining chromosome 8q transcripts of the frozen consensus, plotted on the same colour scale and with the same cohort ordering as Figure 6A. Together the two panels cover the complete 173-transcript cis set. Colour gives the Spearman correlation with PTK2; grey indicates rho <= 0.

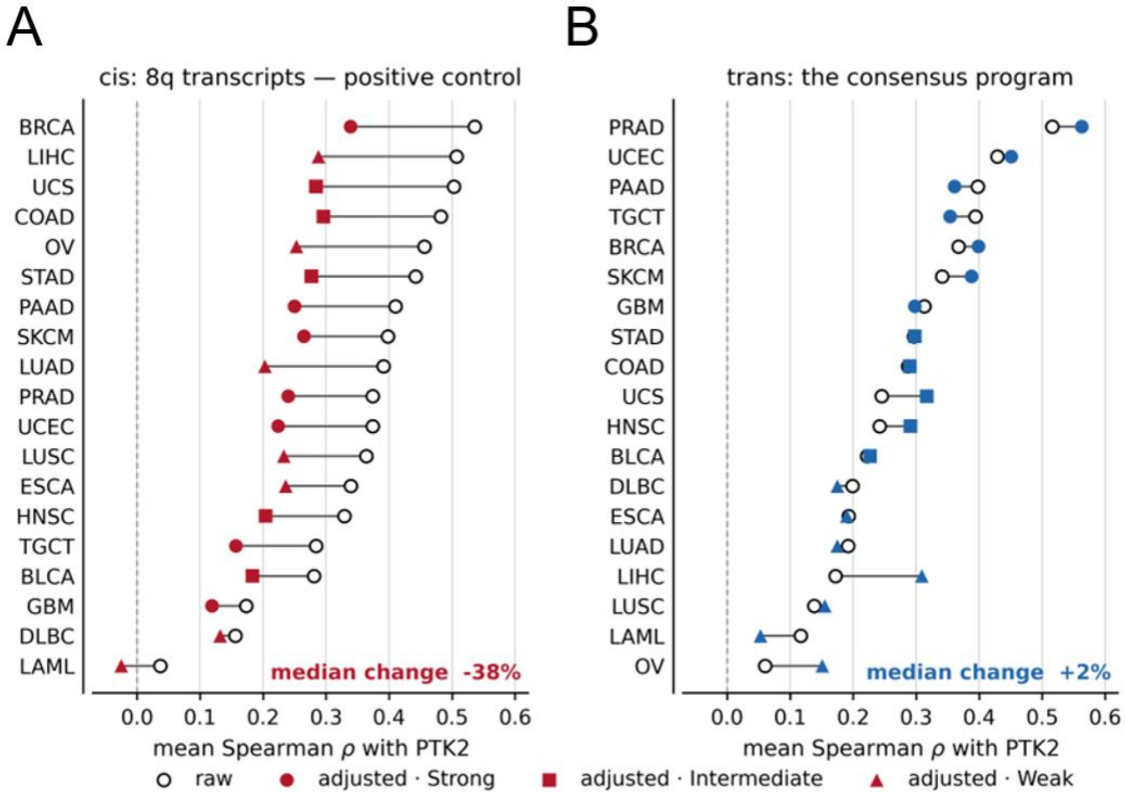


**Figure S4. Co-expression is not explained by copy number.** Mean Spearman correlation between PTK2 and each transcript set by cohort, before (open circles) and after (filled symbols) partial correlation against chromosome 8q arm copy number and genome-wide aneuploidy score. (A) Chromosome 8q (cis) transcripts, which serve as the positive control because they lie on the same arm as PTK2 and must attenuate if the covariate is effective; they fall by 38% on median. (B) The trans consensus, which does not attenuate (median change +2%). Symbol shape denotes the cohort tier assigned in Figure 6C.

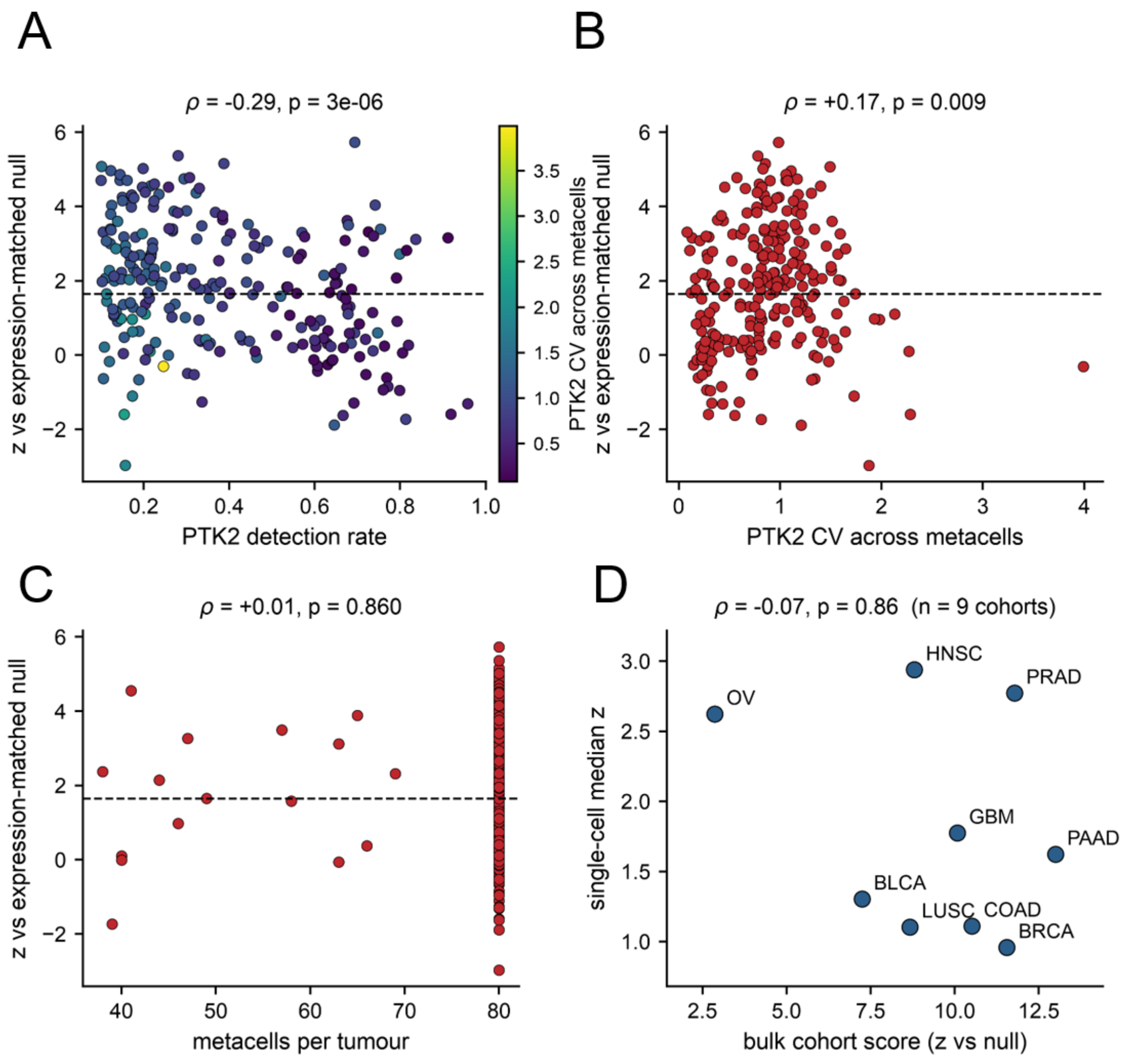


**Figure S5. Single-cell diagnostics and comparison with the bulk analysis.** (A) Co-expression z against the PTK2 detection rate, coloured by the coefficient of variation of PTK2 across metacells. (B) z against that coefficient of variation. (C) z against the number of metacells per tumour. (D) Median single-cell z per tumour type against the corresponding bulk cohort score, for the nine cohorts present in both analyses. Spearman rho and p are given on each panel. The negative association with detection rate and the positive association with the coefficient of variation describe a measurement constraint: where PTK2 is detected in nearly all cells its expression varies little between metacells, leaving little variation against which co-expression can be measured.

### Supplementary Table Legends

The tables themselves are provided in the accompanying workbook Supplementary_Tables.xlsx.

**Table S1. Data sources and software.** Primary data resources and analysis software used throughout, with references.

**Table S2. Gene-set definition parameters.** Every parameter fixed when the gene set was frozen, written directly by the analysis so the table cannot diverge from the code.

**Table S3. The frozen PTK2 co-expression gene set.** Every transcript in the consensus, classified cis (chromosome 8q) or trans, with cytoband, chromosome arm, the number of defining cohorts in which it passed, its mean correlation across cohorts and the current HGNC symbol.

**Table S4. Per-cohort co-expression scores with expression-matched nulls.** Mean Spearman correlation between PTK2 and the frozen sets in each cohort, with the mean and standard deviation of 1,000 expression-decile matched random gene sets, the resulting z-score and empirical p value, and the tertile assignment. Because 1,000 permutations were used the smallest attainable p value is 0.001, so cohorts are compared by z-score.

**Table S5. PTK2 correlation matrix for the frozen gene set.** Spearman correlation between PTK2 and each transcript of the frozen set in every cohort, with no correlation threshold applied.

**Table S6. Partial correlation controlling for copy number.** Mean correlation before and after removing chromosome 8q arm copy number and genome-wide aneuploidy by partial Spearman, with the fractional attenuation for each cohort and transcript class.

**Table S7. Chromosome-arm distribution of the trans programme.** Permutation enrichment across chromosome arms tested against the universe of measured transcripts rather than the whole genome, which controls for gene density. Empirical p values from 10,000 permutations, Benjamini-Hochberg adjusted across arms.

**Table S8. Pathways separating conforming from nonconforming cohorts.** Preranked gene set enrichment analysis per cohort over all measured transcripts, contrasted between tertiles. A term is compared only where it was tested in at least five cohorts of both tiers, and mean normalised enrichment scores are taken over the cohorts in which the term was actually returned.

**Table S9. Preranked GSEA results per cohort.** Normalised enrichment score, nominal p value and FDR q value for every gene set in every cohort, restricted to FDR < 0.25 for size.

**Table S10. Pathway sharing within each tier.** The number of pathways reaching FDR < 0.05 in k cohorts of a tier.

**Table S11. Transcription-factor enrichment.** Enrichr TF Perturbations Followed by Expression tested against the universe of measured transcripts as background, computed separately for the trans consensus and for a size-matched set of PTK2's strongest pan-cancer correlates constructed without the recurrence requirement.

**Table S12. Transcription factors by set membership.** Which factors are enriched in the trans programme only, in PTK2's top correlates only, or in both.

**Table S13. Retired to current gene symbols.** Retired and alias symbols resolved to current HGNC symbols. A symbol was remapped only where the resolved target was unambiguous and was not already present in the expression matrix as a separate row.

**Table S14. Transcripts of the nonconforming cohorts.** For each nonconforming cohort, its most strongly PTK2-correlated transcripts drawn exclusively from genes that failed the recurrence requirement, with the recurrence count of each. These are the transcripts shown in Figure 6D.

**Table S15. Overall survival by PTK2 expression.** Patients divided at the cohort median of PTK2 expression. Hazard ratios compare PTK2-high with PTK2-low by Cox proportional-hazards regression fitted on the same binary split, so that the hazard ratio and the log-rank p value describe a single comparison.

**Table S16. Per-tumour single-cell co-expression results.** For every tumour scored, the number of cells and metacells, the number of trans transcripts above the detection floor, the PTK2 detection rate and coefficient of variation across metacells, the co-expression score, the mean and standard deviation of the within-tumour expression-matched null, the resulting z-score and empirical p value, and the mean correlation for the trans set and for 300 randomly chosen genes. Tumours in which fewer than 20 transcripts cleared the floor were not scored and carry empty statistics.
